# Elephant rumble vocalizations: spectral substructures and superstructures

**DOI:** 10.1101/2024.03.10.584305

**Authors:** Emmanouela Rantsiou

**Affiliations:** School of Engineering and Applied Sciences, Harvard University, Cambridge, MA, USA

## Abstract

Elephant communication, particularly through infrasound rumbles, plays a pivotal role in their social interactions, yet the complexity and functional significance of these vocalizations remain only partially understood. This study explores the spectral characteristics of male African Savannah elephant rumbles, revealing a rich substructure within what has traditionally been viewed as homogeneous low-frequency calls. Our formant frequency analysis of wild male elephant rumble vocalizations demonstrates that rumbles exhibit significant within-call spectral variation, challenging the notion of rumbles as uniform acoustic units. Our findings further reveal that elephant rumbles contain complex patterns of frequency modulation, including distinct vowel-like elements, suggestive of sophisticated vocal tract manipulation. We also document the presence of collectively oscillatory behavior in the formant frequencies of elephant rumbles during group vocalization events. This ‘superstructure’ emerges clearly when silent intervals between rumbles are removed, revealing an oscillatory trend in the chronological sequence of vocalizations. The discovery of intricate spectral substructures and superstructures within elephant rumbles and rumble exchanges respectively, highlights the sophistication of elephant communication systems. This study underlines the elephants’ ability to engage in complex vocal modulation and may have implications for the understanding of their social organization and cognition. Furthermore, unraveling the complexity of elephant vocalizations can enhance our ability to monitor and conserve these majestic creatures, offering new perspectives for studying their social interactions and behaviors in the wild.

## Introduction

Elephants produce a range of different vocalizations including the snort, trumpet, croak, bark, chuff, rev, and various types of rumble calls, covering fundamental frequencies from the infrasound ∼ 10 Hz up to 700 Hz [1–7].

Rumbles - prolonged vocalizations in or near the infrasound range - are reported as being the most frequent calls [2,5,8] and despite the lack of consensus on the number or types of rumbles, their functional roles have been the focus of many studies: it has been shown that rumble calls carry cues on identity [9, 10], reproductive state and sex [11, 12], emotional state and affect [10, 13], but also that they can serve as warning [14] or referential signals [15].

Although rumbles are understood to be produced in the larynx [16], the precise production mechanism was originally debated due to their very low frequency content. The ‘active muscular contraction’ mechanism (the same mechanism believed to drive purring sounds in feline cats [17–19] - but see [20] for the latest update) was suggested as a possible explanation because it allows for arbitrarily low fundamental frequencies [21], independent of the size of the vocal folds. Another explanation, the ‘myoelastic-aerodynamic mechanism’ [22–24] was tested and confirmed, demonstrating that elephant rumbles are produced by flow-driven, self-sustained vocal fold vibrations [21], similar to the mechanism of vocal production in humans and many other animals. Moreover, it has been shown that elephants can selectively engage either the nasal or oral vocal tract for rumble vocalizations, depending on the context [25], but also produce coupled nasal and oral vocalizations [26].

Despite the similarities of the elephant voice production mechanism to that of humans and other animals, the elephant vocal apparatus exhibits noteworthy differences. These include substantial anatomical variations in the vocal folds, such as their positioning and orientation relative to the trachea, and operational differences like the duration of the vibratory cycle, compared to humans [27]. Additionally, the presence of a pharyngeal pouch [28–30] remains a yet unexplored aspect in the elephant vocal production.

In a review of low frequency elephant vocalizations [6], the author mentions the capability of elephants to lengthen their vocal tract not only by extending their trunk, but also possibly through the descent of their larynx. In terms of trunk extensibility, a study focusing on elephant trunk bio-mechanics [31] found a significant elongation capacity of 135% (ratio between longest and shortest states), although this observation was not specifically linked to vocalization processes. Concerning the lowering of their larynx, it is proposed that the unique anatomical structure of the elephant hyoid apparatus and associated muscles imparts considerable laryngeal mobility, facilitating the generation of infrasounds [16, 29].

An increasing list of studies are showcasing the ability of various species to modify their vocal tracts [15, 32–44]. In the case of some primates in particular, vocal tract manipulation has been shown to include articulatory movements such as changes in position of the lips, the larynx, and the jaw [34, 36, 40, 41, 43]. This active vocal filtering results in formant (i.e. vocal tract resonance frequencies) manipulation and phonetically contrasting sounds, akin to human vowels. Indeed, the presence of vowels or vowel-like spectral structures is explicitly mentioned in several primate and non-primate studies [35, 38, 40–46].

Published studies on elephant vocalizations analyse rumbles as vocal units: spectral features are extracted as averages over the whole vocalization or over one pre-selected section - typically in the middle - of the rumble (see e.g., [9, 13, 25, 47, 48]). There is a good reason for that: rumble vocalizations are continuous sounds, that exhibit flat or minimally and progressively varying fundamental frequencies. Based on those spectral and structural characteristics, a rumble could be seen as an analog to a note, the basic acoustic unit in vocalizations or communication systems of various species including birds and primates. However, as our study will demonstrate, rumbles exhibit spectral substructures and variations. Investigating these substructures within rumbles is crucial for identifying the basic acoustic units in elephant vocalizations - akin to whale clicks [44, 49–52] or human phonemes [53–55] - and for exploring their potential functionality or their associated semantic information.

In this study, we analyze formant frequencies in order to examine the vocal structures and modulations in elephant rumble vocalizations and the implications for articulatory movements in elephants. Our findings suggest that elephants engage in active vocal filtering, shifting their formants not only between, but also within rumbles. In the ‘Results’ section, after presenting evidence that the rumbles in our dataset showcase internal substructure, we investigate the implications of such structures on vocal tract manipulation and suggest that rumbles can have vowel-like qualities. We map the vocal rumble space of the elephants in our dataset and attempt to make a qualitative comparison to known vowel spaces of primates. We finally make an intriguing observation: the formant frequencies (and in particular the second formant F2) of the rumbles in an antiphonal exchange, build collectively, rumble by rumble, an oscillatory and at times periodic pattern. Separate ‘Methods’ and ‘Supporting Information’ sections include technical details, additional and clarifying plots, as well as a link to a repository with the reduced data and other relevant information.

## Data

The data analyzed in this study consist of a reduced dataset of rumble vocalizations, previously used in [56]. These vocalizations took place in the wild within male elephant groups, in the context of group visits to a waterhole area. The rumbles are antiphonal vocal exchanges involving two or more members of a male elephant group and serve to communicate and coordinate the groups collective departure from the watering ground. Each waterhole visit typically includes three to four phases: 1) an initial silent period with no vocalizations, 2) a - not always present - sequence of rumbles from a single caller (the dominant elephant), 3) a sequence of rumble bouts where two or more elephants - but not necessarily all of the group’s elephants - engage in antiphonal exchange of vocalizations, and 4) the resulting collective retreat of the elephant group from the waterhole. We refer to each one these group vocal exchanges as an ‘event’. The reduced dataset used in this study includes rumbles from six such events. The number of rumbles as well as the number and identity of group members vary among these events. 116 rumbles from 13 elephants are used in this study (Table 1). Some individuals appear in more than one of these group vocal exchanges. It is important to mention that spectral features were not extracted as averages over the whole duration of a rumble, as it is usually done in the literature. Instead, each rumble was segmented in consecutive time slices of 0.25s, and formants F1 and F2 were extracted for each such slice. Formants F3 and F4 where not always present in the rumbles, but when available we extracted them as averages over the full rumble duration. Further information, as well as the reduced data used here can be found online as part of a previous publication [56] and in the ‘Supporting Information’ section of this manuscript.

**Table 1:** Summary of the data used. 6 rumble exchange events, each with different group composition and size, and number of rumbles exchanged.

| **Event ID** | 05 | 07a | 07b | 11a | 11b | 17 |
| --- | --- | --- | --- | --- | --- | --- |
| **# of callers** | 4 | 5 | 3 | 3 | 5 | 4 |
| **# of rumbles** | 21 | 26 | 8 | 11 | 30 | 20 |

## Results

In the section that follows we will present analysis that support three findings:

1. rumbles have internal spectral structure,
2. vocal tract manipulation is required to produce these structures, which show vowel-like qualities,
3. antiphonal rumble exchange events show collectively oscillatory behaviour.

### Spectral sub-structure of rumbles

Typically, when examining the spectral content of rumble vocalizations, a single rumble call is considered an audio unit and any feature extraction, like fundamental or formant frequencies are calculated as averages over the whole duration of the rumble. However, neither the fundamental nor the formant frequencies, necessarily remain fixed within a rumble call. On the contrary, as we will show, spectral content variations within rumbles can be substantial. Importantly, this fact has direct implication for efforts to understand the vocal communication code of elephants.

Fig 1 shows the range of F1 and F2 values for all rumbles in our dataset. Formants are calculated either as averages over full rumbles (left panel), or from slices of 0.25s (right panel), as previously mentioned. We notice that when refining the time-resolution over which we calculate formant frequencies, the range of formants, in particular that of F2, increase considerably - from 15-34Hz to 8-46Hz for F1 and from 65-94Hz to 56-105Hz for F2 - indicating that when averaging over the full rumble duration, we overlook the ‘sub-rumble’ structure of these spectral features.

**Figure 1:**
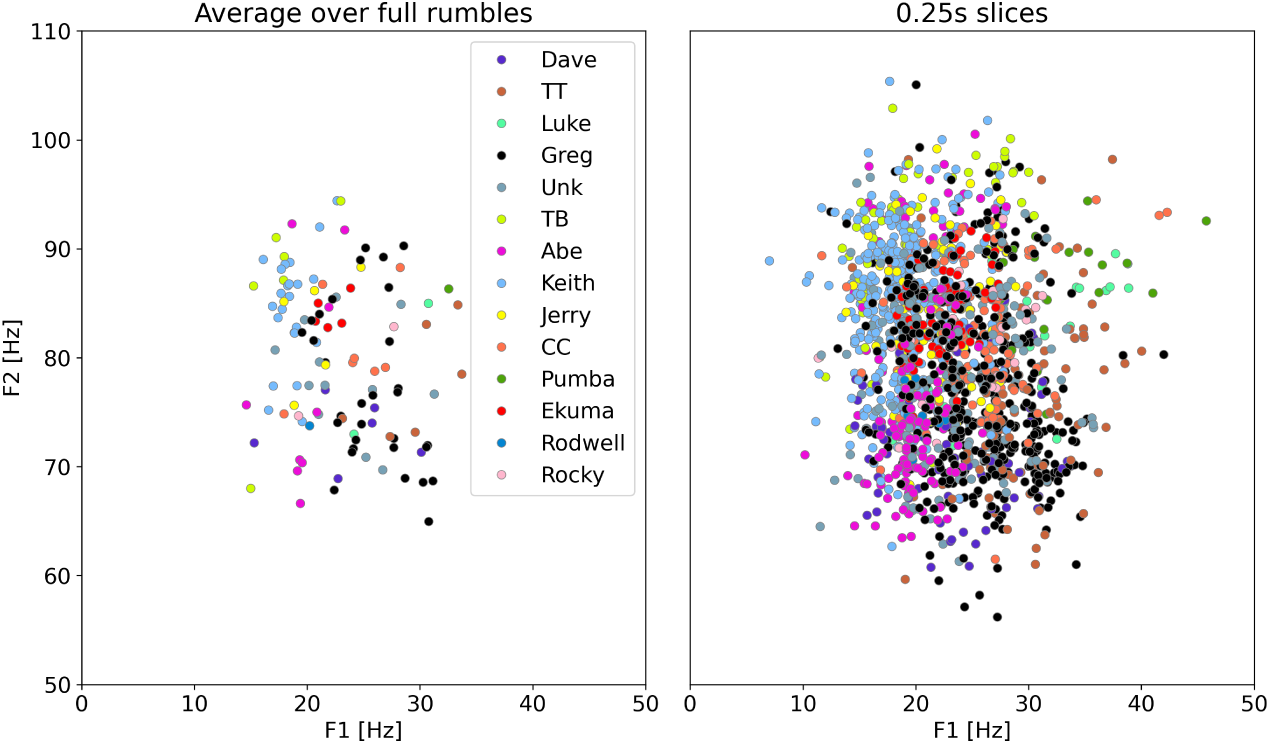
Range of formant frequencies. Averaging of formant frequencies underestimates the range of F1 and F2 of rumble vocalizations and misses the variability of spectral features within single rumbles. Formant averages over full rumbles result in ranges of 15-34Hz and 65-94Hz for F1 and F2 respectively. Slicing up rumbles to 0.25s segments gives value ranges of 8-46Hz and 56-105Hz for the first and second formant respectively.

In Fig 2 we show the formant frequency ranges (for rumble vocalizations) of nine elephants. These are individuals for whom we have more than five rumbles in our dataset. All rumbles were divided into slices of 0.25s and formants were extracted for each such slice, shown in the plot as individuals points overlaying the boxplots. Here we note that the range - and distribution - of formant frequencies vary among individual elephants, as discussed in [56]. Additionally, formant ranges also vary from event to event, as can be seen in S2 Fig of the Supporting Information.

**Figure 2:**
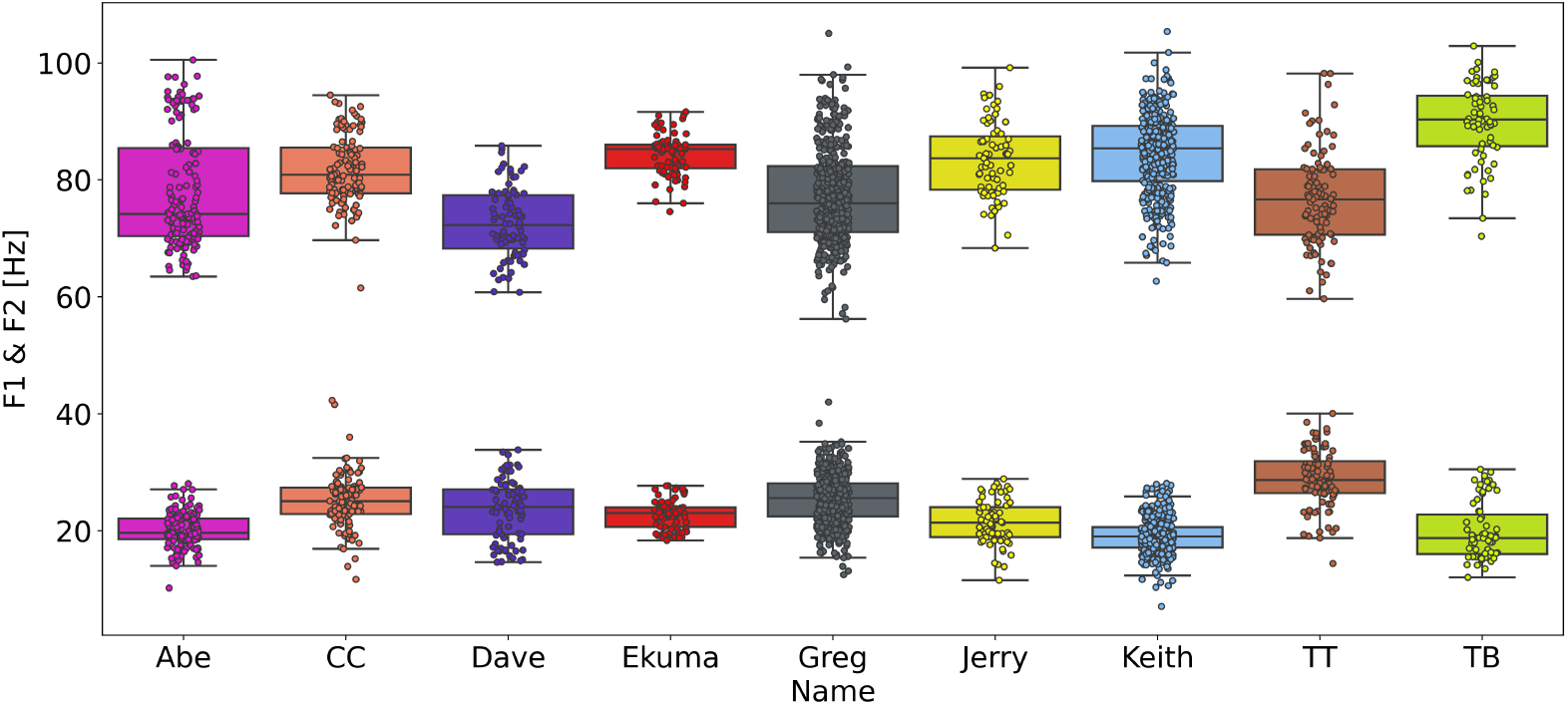
Range of rumble formant frequencies for individual elephants. Each pairs of boxplots describe the F1 and F2 ranges of each elephant in our study. All available rumbles for each individual are segmented in 0.25 of non-overlapping slices. Formant frequencies F1 and F2 are extracted from each of the slices. Additional stripplots are overlaid on the boxplots, to better visualize the density distribution of the formant values.

Fig 3 shows variations of F1 and F2 within single rumbles for 24 rumbles from three different elephants. F1 shows, in general, smaller relative variation than F2, and we notice rumbles with significant inter-rumble formant variation and others with almost flat F1 and F2 profiles throughout vocalization. This latter observation is illuminated further in Fig 4. Four rumble spectrograms are shown together with the corresponding F2 values calculated on 0.25s slices of the original rumble. Two of the spectrograms show noticeable variability in F2 values, while the other two have a slight downward trajectory (bottom right) or flat (bottom left) formant profile throughout the duration of the rumble. Although rumbles are continuous sounds, i.e., they lack any pauses (an intensity drop to or near zero), the above clearly show their spectral content can be variable. In S3 Fig we show the narrowband spectrograms of the same four rumbles together with their spectra (0-200Hz) and the F1 and F2 peaks detected with LPC, this time calculated over the full rumble.

**Figure 3:**
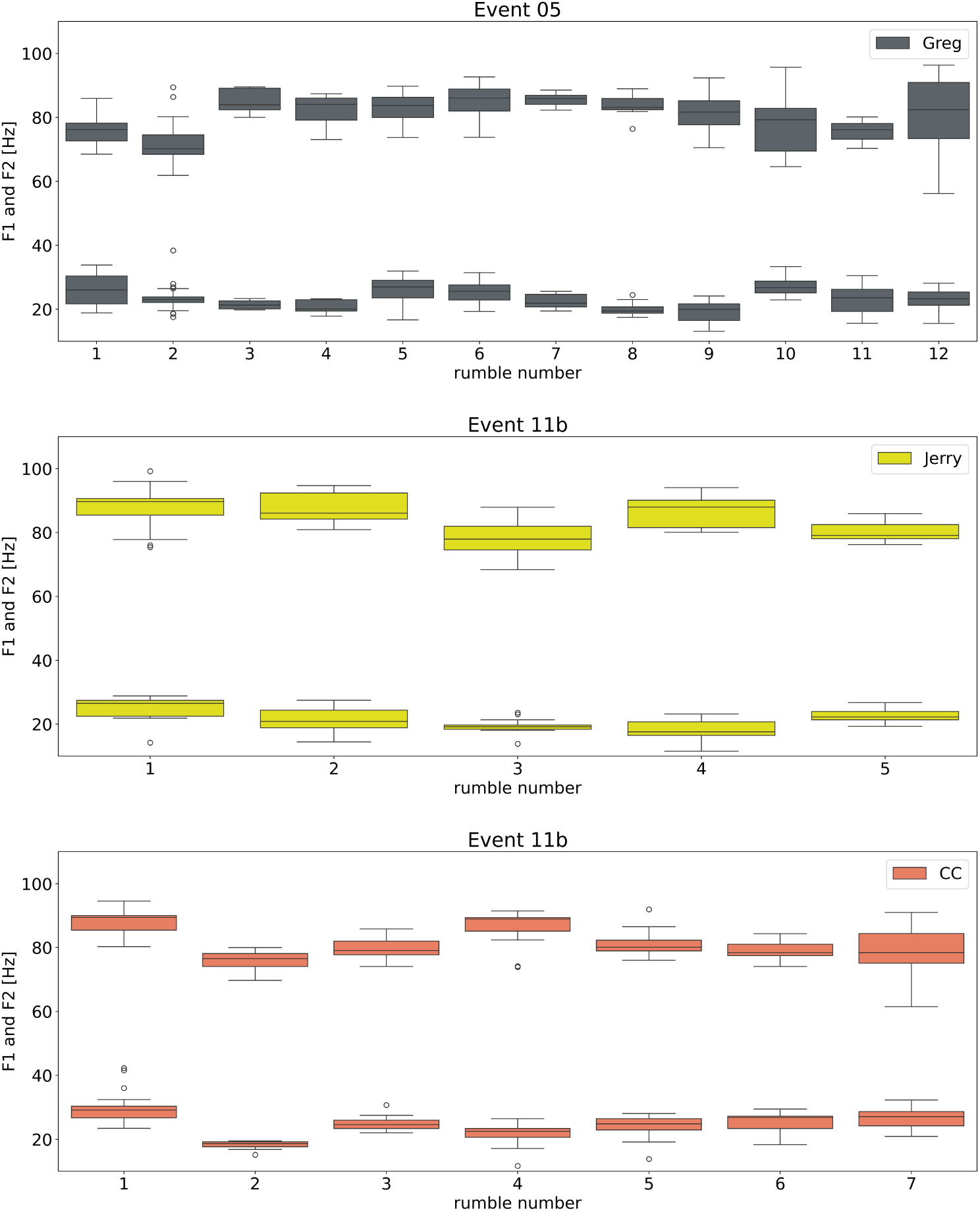
Range of formant frequencies within single rumbles. Each pair of boxplots indicates the variation of F1 and F2 in a single rumble. The three plots correspond to three elephants and each of them include all the rumbles that elephant produced in an event. Each rumble was segmented in 0.25s non-overlapping slices and the formants were calculated on each of these slices.

**Figure 4:**
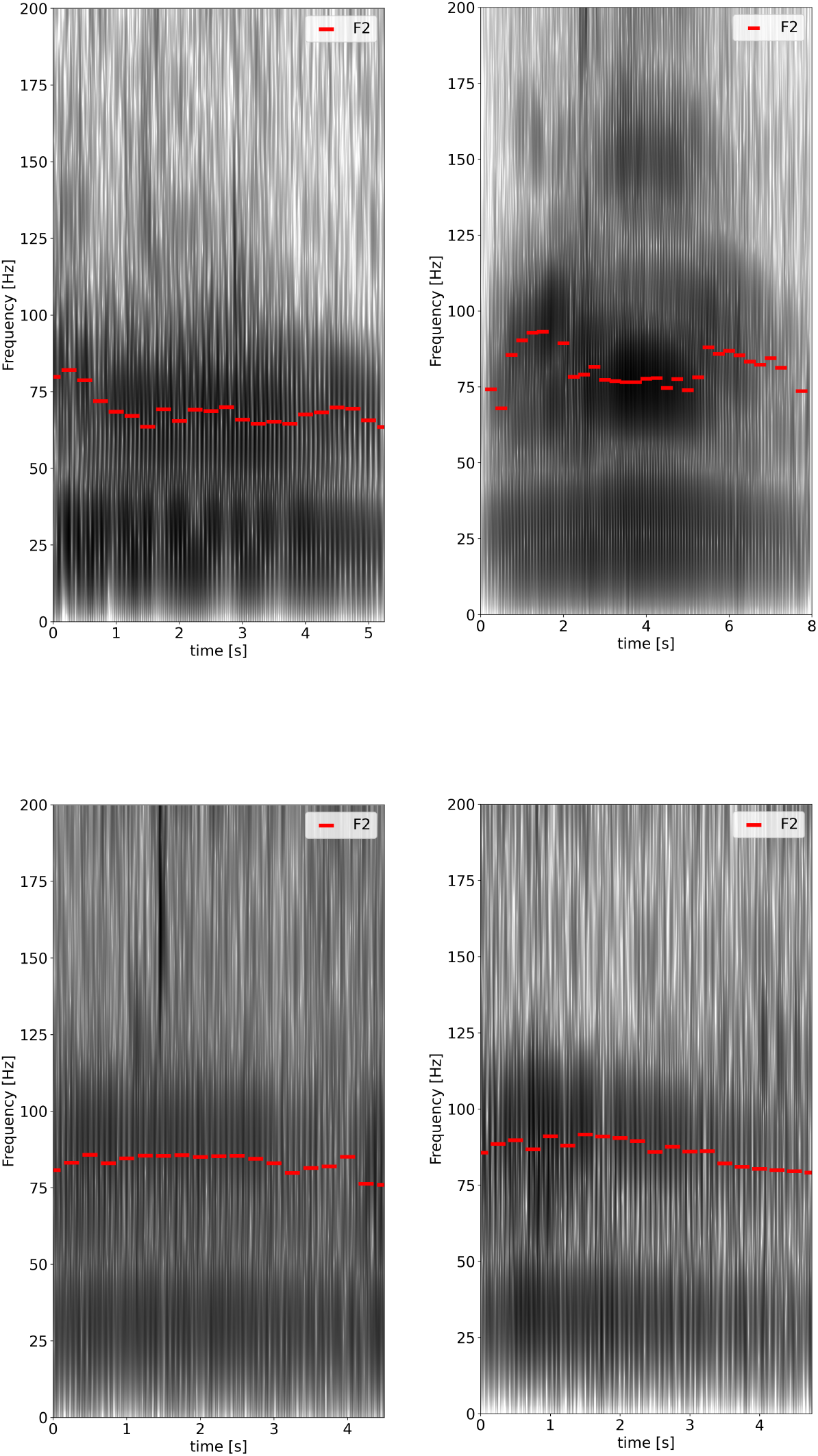
F2 variation within single rumbles. The formant frequencies within a rumble call may vary considerably. The spectrograms of four different rumbles are shown here with the F2 values of each rumble shown as red dashes. The formant values are calculated for windows of 0.25s from the audio files. The top figures showcase clear formant variation while the bottom ones have rather flat and steady (left) or slightly downwards (right) F2 frequency trajectories throughout the duration of the rumble.

### Vowel-like patterns in rumble vocalizations

The variation of the formant frequencies within and across rumbles as described above, brings about interesting questions and observations regarding voice production and Vocal Tract (VT) manipulation in elephants.

In the uniform tube model of voice production, the vocal tract is approximated by a tube of uniform cross-sectional area closed at one end (the vocal folds) and open at the other (the mouth or nasal end). This is a quarter-wave resonator which can sustain standing waves of integer multiples of 1/4 the source wavelength - the source here being the vibration frequency of the vocal folds. The relation between the resonator’s length *L* and the resonant frequencies (formants) it can sustain is:

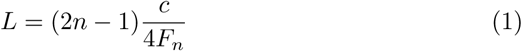

where c is the speed of sound in air, and *F_n_* the n-th formant frequency in Hz (see any physics textbook; e.g., [57,58]). Böe *et al.* [59] observed that uniformity (sounds produced from a uniform tube) would inevitably result in agreement of vocal tract length (VTL) estimations from all formants when using Eq. 1, whereas deviation from that would be indicative of a non-uniform tube. This has been shown for humans [59] and baboons [35]. We used formants F1 to F4 to estimate the VTL according to Eq. 1. As shown in Fig 5, the distribution of values vary significantly, depending on the formant values used. Formants F1 and F2 give mean estimated values of 3.9 m and 3.2 m respectively, while F3 and F4 agree to a mean VTL value of 2.2 m. The discrepancy between the estimated lengths is significant, suggesting that the assumption of uniformity is violated. Additionally, the limit values of the derived VTLs - 1.2 m and ∼ 6 m are anatomically implausible.

**Figure 5:**
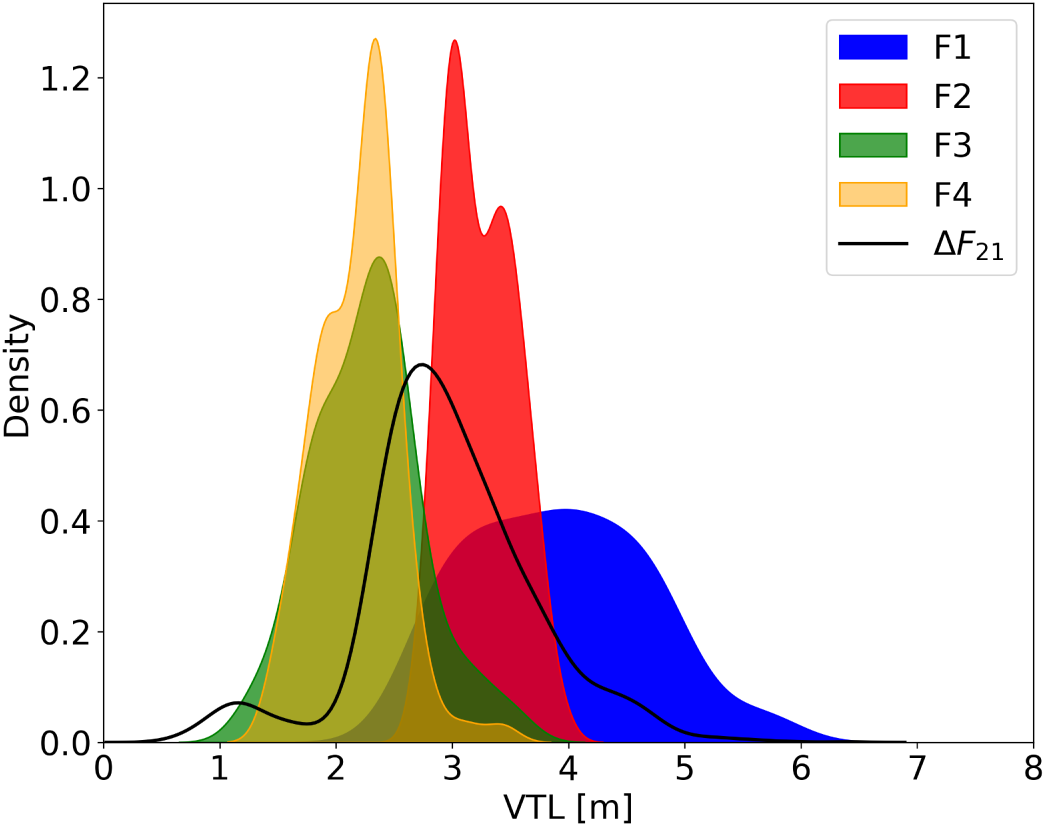
Vocal Tract Length estimated by formants. Distributions (Kernel Density Estimations) of VTL based on the first four formant frequencies (in colors) and the formant dispersion F2-F1 (black line), assuming the fixed-tube model.

An alternate and more robust way to estimate VTL in cases of non-uniform tube configurations has been suggested and demonstrated by Fitch [60], and relies on using the formant dispersion, i.e., distance of successive formants, as opposed to formant values (for application of formant dispersion on elephant vocalizations see e.g., [9, 47]). Using the formant dispersion Δ*F*_21_ of the first two formants resulted in an anatomically realistic mean VTL of 3.1 m.

Values of elephant VTLs reported in the literature are approximate and there is a lack of systematic documentation of VTLs for Loxodonta africana in general. Soltis [7], using reports from measurements of mandible [61] and trunk lengths [30] [62], estimated that the total VTL for an adult female L. africana is 2.5 m (0.75 m for the oral VT and another ∼ 1.8 m for the trunk length). Male African savannah elephants are significantly larger than females, with typical trunk length for adults of *>* 2 m, although published reports or measured lengths are sparse. We adopt a VTL of 3.0 m for the discussion that follows.

Comparative studies of animal vocalizations and in particular of primates, often superimpose the F1-F2 plane of human vowels to that of other animal vocalizations. We attempt to compare the formant space of elephant rumbles to the vowel space of human and other primate vowel-like vocalizations. Scaling or normalization of the frequencies is required due to differences in VTLs among species [59,63,64]. Here we present a comparison of the formant space of elephant rumble vocalizations and the vowel space of humans (Fig 6), as well as that of non-human primates. To do so we scale the rumble formant frequencies to the VTL of humans using the relation:

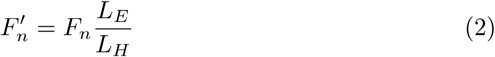

where *L_E_/L_H_* is the ratio of elephant to human VTLs, and 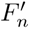 is the scaled n-th formant. The human vowel space data and relevant VTL values come from Peterson and Barney [65] and are publicly available.

**Figure 6:**
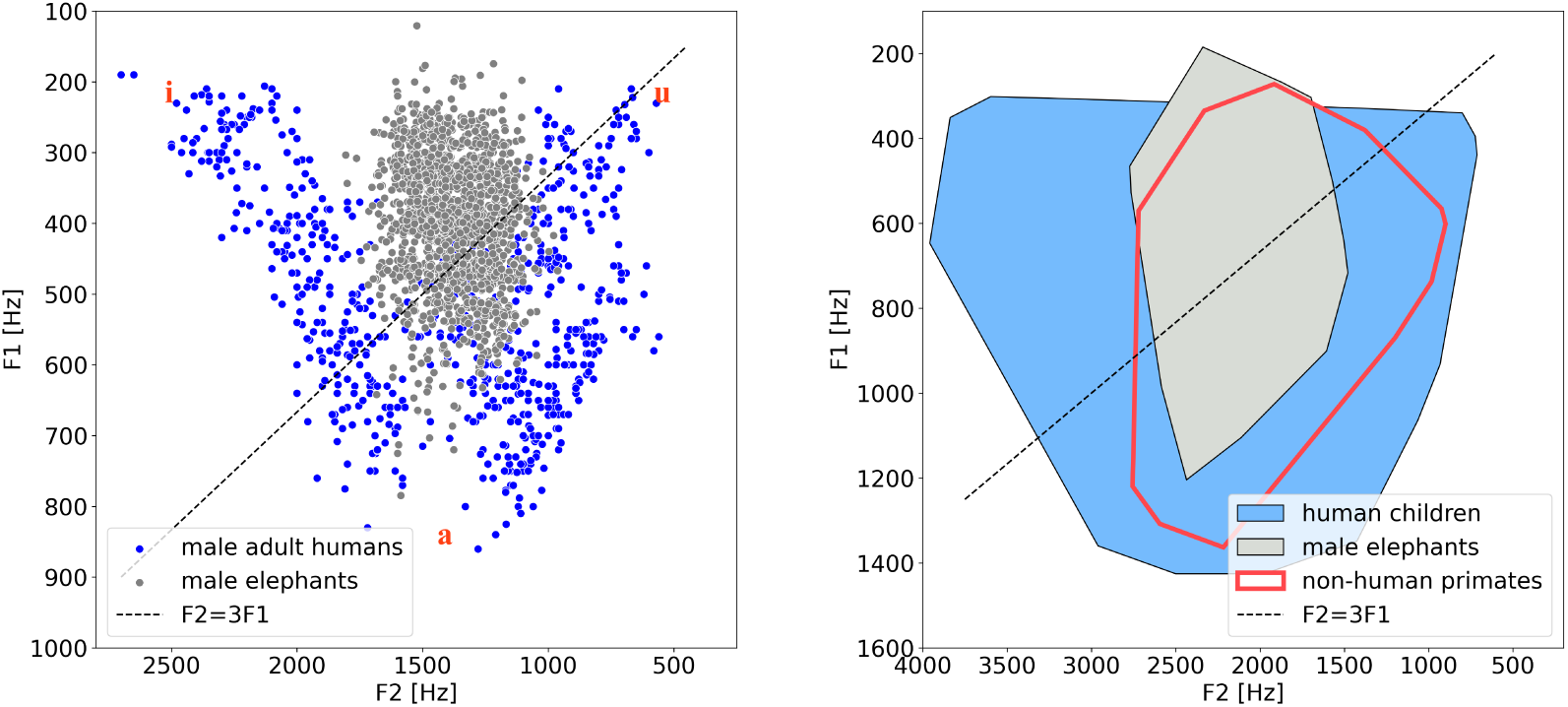
Comparison between the vowel spaces of primates to the vocal (rumble) space of elephants. *Left:* the formant space of all elephants (gray dots) in our dataset (rumble calls) is compared to the vowel space of human adult males (blue dots). The elephant formant frequencies were scaled to the average VTL of male adult humans (17.5 cm). *Right:* Convex hulls of human children vowel space (blue), male L. africana rumble formant space (gray), and non-human primate vowel space (red). All data were scaled to a VTL=11.4cm (corresponding to the VTL of a macaque in [40]). The collective vowel space for the non-human primates is from Fig 2 in [59], and compiles studies available up to the time of that paper’s publication. It includes vocalizations from Diana monkeys, mongoose lemurs, chacma baboons, hamadryas baboons, gorillas, and rhesus macaques (see paper for list of source references). The black dashed line in both figures corresponds to *F* 2 = 3*F* 1, which gives the expected relation between the first two formant frequencies in the uniform tube model. The data for humans are from Peterson and Barney [65]. Note that the axis assignment and direction of increasing values (leftwards and downward) follow the convention used in the phonetics literature.

The intention of the comparison presented in Fig 6 is to 1) show that elephant rumble vocalizations do not follow the uniform tube model, but require VT manipulation, and 2) compare the range of rumble vocal space to that of the vowel space of humans and other primates, in order to get an estimate of the relative extent of the vocal space of elephants (for rumbles only).

### Collectively oscillatory behaviour of rumbles

In five out of the six group vocalization events examined in this study, we have noticed the presence of rumble ‘superstructures’: a collectively oscillatory (and potentially periodic) behaviour of the formant values of the analysed rumbles, and in particular of the second formant F2. This becomes apparent - as demonstrated in Fig 7 - after removing the silent time between rumbles and plotting the first two formants of the rumbles in the order in which the vocalizations took place.

**Figure 7:**
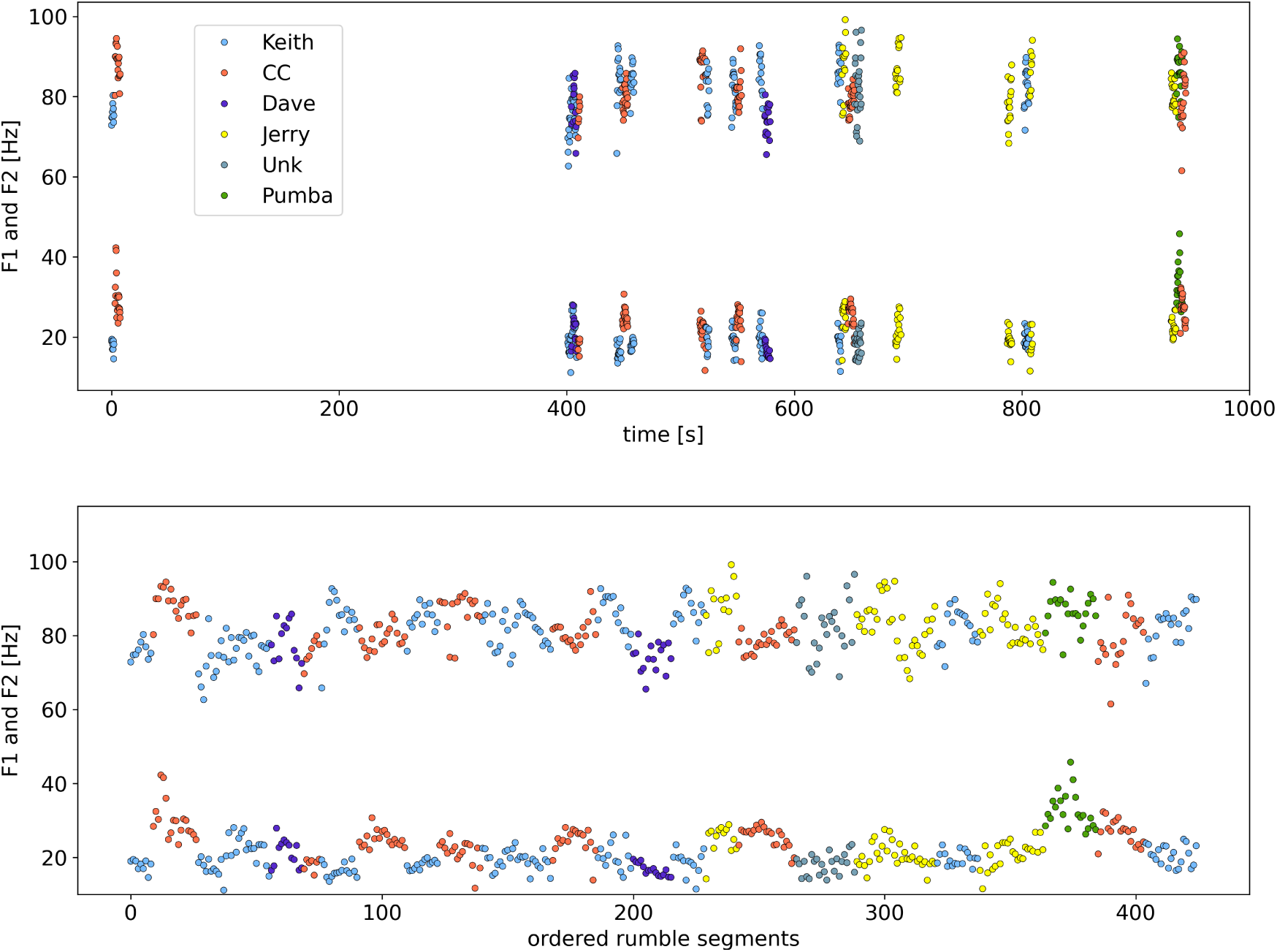
Oscillatory trajectory of formants in rumble exchange event 11b. *Top:* The formants F1 and F2 of all rumbles in event 11b are plotted, retaining the chronological order of utterance of the rumbles. Each rumble was segmented in slices of 0.25s and each such slice is represented by a point. Color coding identifies the vocalizers. *Bottom:* Removing the inter-rumble time (silences) and plotting the formant progression of each rumble consecutively in the order the rumbles took place, we notice an oscillatory behaviour.

Fig 7 shows the trajectories of the formant frequencies F1 and F2 of all rumbles in event 11b. Each point on the plot corresponds to a 0.25s segment of a rumble. The top panel shows the progression of the rumble exchange in real time units on the x-axis. We notice five elephants taking turns rumbling for about 16 minutes. We furthermore notice the variation of the formant frequencies within single rumbles but also among different callers. Removing the silent parts in-between rumbles results in the plot in the bottom panel: an oscillatory behaviour of both formants is revealed. Fig. 8 shows three more examples (events) and the relevant figures for the remaining two events can be found in ‘Supporting Information’.

**Figure 8:**
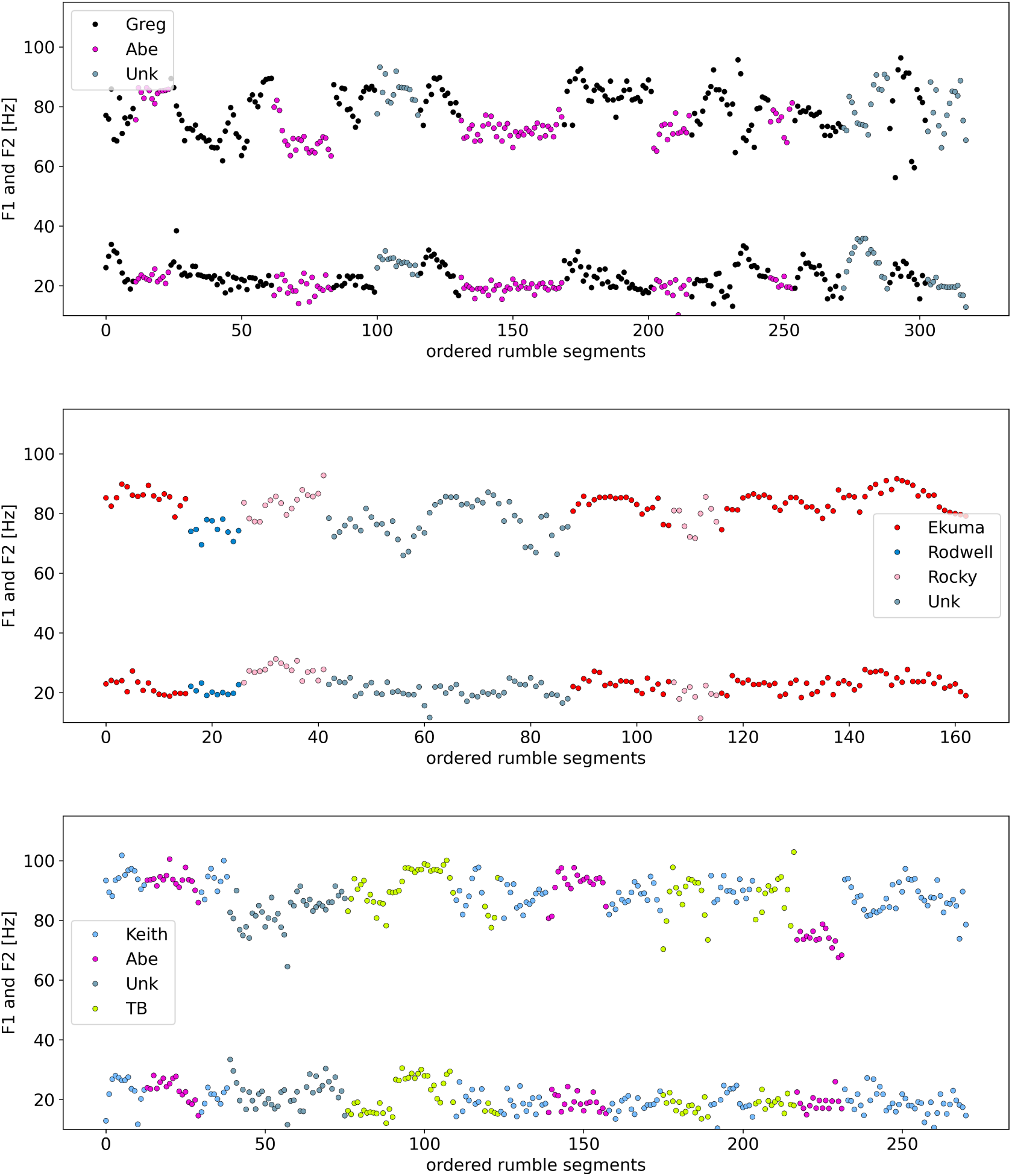
Oscillatory trajectory of formants in rumble exchange events 05, 11a, and 15. Similarly to Fig 7, events 05 (top), 11a (middle), and 15 (bottom) showcase an oscillatory behaviour of formant F2 in particular, visible when the inter-rumble time is removed.

This oscillatory behaviour is not simply a binary high-low, alternating sequence of formants from one rumble to the next, but is put together by the varying formant trajectories within the individual rumbles that comprise the sequence of vocalizations. Together these trajectories form the oscillatory patterns observed in Figs 7, 8. Thus, it appears as if elephants partake in collectively creating, rumble by rumble, a longer oscillatory signal.

We found the presence of these patterns intriguing and felt compelled to attempt a data fit. For two of the events we used symbolic regression (with PySR [66]) to find mathematical expressions that could best describe the patterns observed. We focused only on F2 as the pattern manifests more strongly there than in formant F1. Fig 9 shows fit curves for events 11a and 15. In both cases the fit is a superposition of sinusoidal functions and describes, at least in a qualitative way, the oscillatory trends of the data. Details on the fit parameters and choice of functions can be found in ‘Methods’.

**Figure 9:**
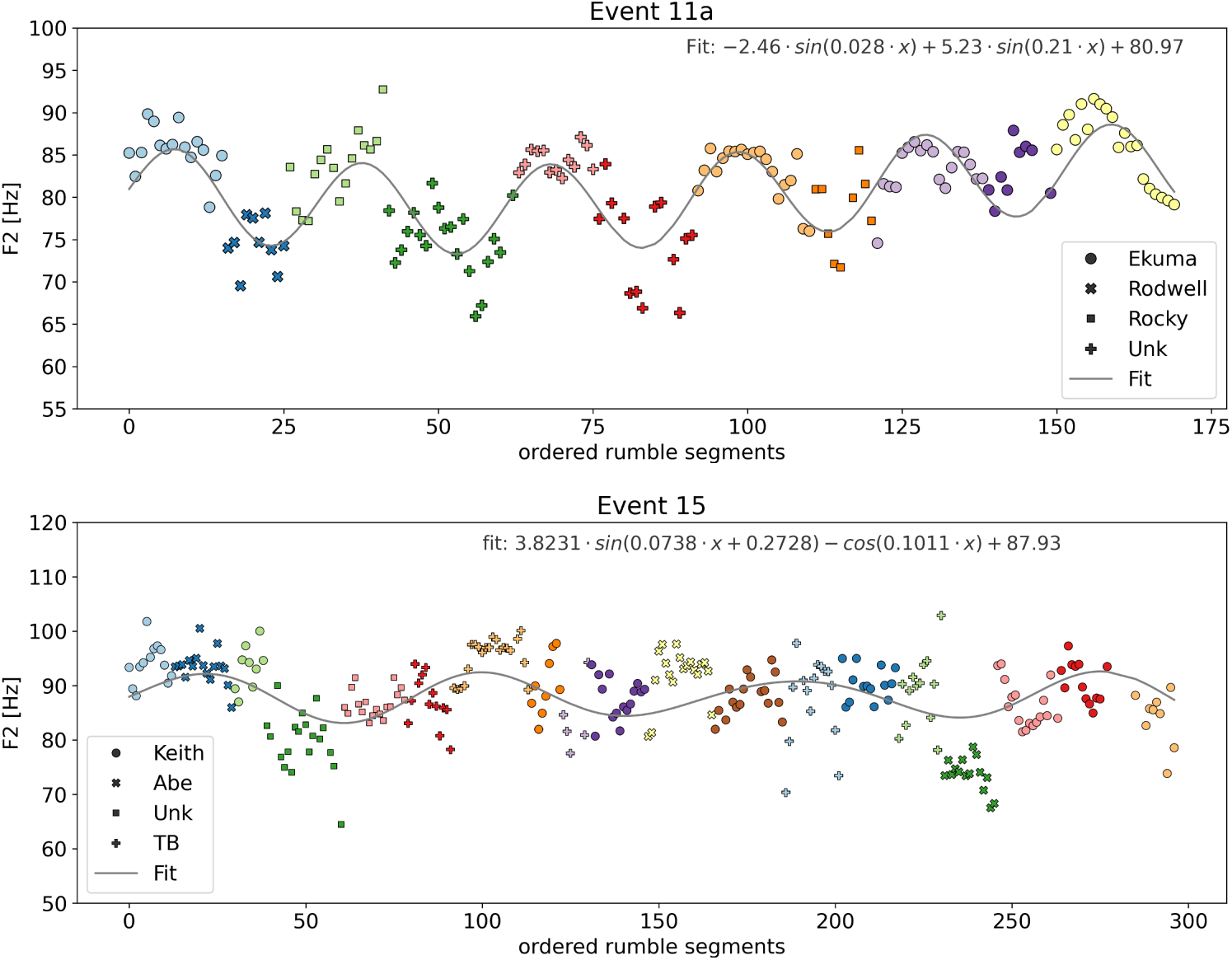
Fit curves on oscillating formant F2: events 11a and 15. Fit of F2 for two rumble exchange events. The alternating colors indicate transition from one rumble to the next. The identity of each caller is indicated by its own symbol. Both events can be fitted with waveform functions resulting from the sum of two sinusoidals and an offset. Residuals and further plots can be found in the Supporting Information section.

Residual plots included in the ‘Supporting Information’ section show the quality of the fits. We observe local or more generalized failures to fit amplitude or phase, made obvious by the fact that residuals follow the pattern of the original data. However, the purpose of the fit curves is to serve as visual aids in describing the oscillatory patterns observed.

The role of these oscillations in the functionality of antiphonal rumble exchanges as pre-departure communication among elephants remains, for the moment, unknown to us. We attempted however, to rearrange the order of vocalizations (rumbles) in these antiphonal events to examine how these oscillatory patterns would be affected. In ‘Supplemetary Information’, we present plots where we “scrambled” the order of rumbles in various ways to observe the result this would have on the oscillatory pattern. The results are interesting and we include a small discussion to go along with the plots. The main observation that emerges is that randomizing the order of segments within rumbles diffuses the oscillatory pattern observed in the original vocal exchange, demonstrating that these oscillatory patterns did not come about by chance.

## Discussion

In this paper, we demonstrated for the first time that the rumble calls of L. africana exhibit a complex internal spectral structure, characterized by significant variations in the first and second formant frequencies (F1 and F2). These variations resemble vowel-like spectral structures that necessitate articulatory movements from the elephants, comparable to the vocal tract manipulations humans use to produce vowels. While we are presently unable to assign a definitive acoustic function to these variations or make any assumption as to whether they carry semantic information, we propose that their production mechanism is fundamentally similar to the creation of vowels in human speech.

Our comparison of the elephant’s rumble formant space to that of primate vowel space attempted to showcase the relative range of vocalic abilities and articulation potential of L. africana. We note again that in this study we focused only on low frequency rumble vocalizations, which are but one type of the many calls produced by elephants. An extended study, which would include the full repertoire of elephant vocalizations could illuminate the presence, variety, and level of complexity of vowel-like patterns in elephant calls and assist in determining whether they serve a communicative function, but also for comparative linguistic studies to other species.

We also documented the presence of a collectively oscillatory behaviour of the formant frequencies (F1 and F2) in sequences of antiphonal rumble exchanges. To the best of our knowledge, this has not been reported elsewhere in the scientific literature. More specifically, the formant trajectories (in particular of F2) of the rumbles in the antiphonal exchanges examined in this study, form together, when the inter-rumble silences are removed, an oscillatory pattern, which persists either in the entirety of the event, or manifests locally within certain segments of the exchange. In instances where the oscillation is localized, it is plausible that the exclusion of rumbles from one or more individuals could reveal a more pronounced oscillatory pattern, suggesting that those individuals did not contribute to the formation of the vocal structure observed. The data fit attempted for a couple of the vocal exchange events was aimed at capturing the general behavior of these vocal superstructures rather than providing a detailed mathematical model. Although better fitting functions could be achieved, by allowing for more complex combinations of sinusoidal or other functions (see for example an alternative fit for event 15 in ‘Supporting Information’), this would surpass the purpose of this investigation and would currently offer no further insight on the functionality of these oscillations. It is important to note that, without resolving the rumble substructures, we wouldn’t be able to identify the oscillatory superstructures.

We do not yet know the function of these collective oscillations. Our objective in this study was to report, in a qualitative manner, a finding which might have implications on the understanding of the structure of the elephants’ communication system and the semantic functionality of antiphonal rumble exchanges. However, we feel compelled to make an analogy. Given the observation offered earlier, that rumbles can be characterised as notes, and adopting the definition of a ‘cantus’ (a ‘song’ in the strict sense - sensu stricto) from W. H. Thorpe (as seen in [67]): “a series of notes, generally of more than one type, uttered in succession, and so related as to form a recognizable pattern or sequence in time”, we are tempted to name these collectively built oscillatory superstructures as ‘resonance cantus’ or ‘formant cantus’ (or, colloquially, ‘formant song’).

## Methods

### Data Analysis

We used Praat [68], a free software package for speech analysis, for feature extraction. Praat’s scripting capability was used to automate and speed up the procedure. To obtain accurate results, specific parameters must be set beyond the softwares defaults and verified meticulously using a substantial number of vocalization data. This necessity applies broadly to various types of non-human animal vocalizations. The procedure employed to ensure reliable formant frequency extraction using Praat is detailed below. The scripting (via Praat) of the process described below was done for the purpose of reproducibility and automating this procedure for handling large number of files. Importantly, this process allows us to use the same parameter settings for all file treatment, therefore minimizing human bias.

- For formant frequencies extraction:

1. Each audio file is resampled to 1000Hz, giving a Nyquist frequency of 500Hz. Reasoning: visual inspection of all 120 audio files have shown that there is no power above 500Hz (in reality, from a little below 500Hz), even for rumbles that clearly show 3rd and 4th formants. Note: the Precision parameter used for the interpolation during resampling is set to 50 samples.
2. The spectrum of the resampled audio file is calculated (zero padding for Fast Fourier Transform selected)
3. LPC (linear predictive coding) smoothing on the spectrum from the previous step, using 9 peaks and pre-emphasis from 50 Hz.
4. Formant peaks are then identified by Praat from the above smoothed spectrum, with the maximum number of possible peaks set to 5.

The parameters chosen for this process (e.g., the number of peaks for LPC, pre-emphasis limit, etc) were optimized through careful, manual trials to ensure the resulting formant values match those extracted via visual examination of the spectra from approximately 20 audio files. Once the reliability of these parameters was established, the procedure steps described above were scripted in Praat and applied to all audio files (about 120 in total). Scripting allows control over a larger number of parameters than does the GUI, which is adapted to human vocalizations. The resulting formants and fundamentals for each file were then validated through visual inspection of each spectrum. This same procedure, on the same files, was repeated a couple more times: Praat script runs on all audio files, automatically extracting spectral parameters for each one and then those values are confirmed manually (visual inspection) by e.g., identifying the formant bumps in the spectrum for each of the 120 files.

We also used Praat for visual inspection of spectrograms and spectrogram plots. All other audio and statistical analysis as well as plots were done with Python.

We used PySR [66] - an open source package for symbolic regression - to calculate the fits presented in §. We used a limited group of operators (+, −, *, cos, sin, inv) and explicitly discouraged nested trigonometric functions. The reason for the relatively limited and mathematically simplistic type of operators is that our goal was to investigate if a simple (semi-) periodic function could capture the observed patterns. Any too involved fitting function, regardless of its goodness of fit, would defy physical or meaningful interpretation in this particular application. L2-norm was used as the element-wise loss, and for each fit we typically used 30-80 populations and 200-300 iterations. The size of each population and number of mutations were kept to their default vales (33 and 550 respectively). Running on an Apple M1 Pro on 10 CPUs, required on average 15s for one fit. At the end of each run, we choose from the proposed solutions either the one with the lowest loss or the one with the highest score (which is a compromise between low loss and small complexity of the proposed function).

## Acknowledgements

The author would like to thank L. Mahadevan for hosting her in the SoftMath Lab during the preparation of the work included in this manuscript. The raw data of elephant rumbles, the reduced content of which was used for this study, were provided by Caitlin O’Connell. Thanks goes to the Image Analysis Collaboratory at HMS for recommending software solutions for symbolic regression and to Simon F. Nørrelykke for useful comments on this manuscript.

## Supporting Information

**S1 Data** The reduced data used for this study - comprising acoustic features such as formant and fundamental frequencies - can be accessed at the following repository:

https://zenodo.org/records/11489682

For our analysis, we sliced each raw audio file (rumble) in non-overlapping segments of 0.25s. We choose this level of time resolution because we wanted to have as fine time-step as possible while simultaneously being able to have enough cycles (signal) in each segment to correctly identify low frequency content (≥ 10 Hz). In cases where higher formants (F3 and F4) were identifiable, we calculated the average formant values across the entire duration of the rumble, rather than from finely segmented slices. The latter, requires automating this process through Praat’s scripting capability for managing the extensive number of audio segments. However, the relatively lower signal-to-noise ratio for these higher formants makes accurately identifying the F3 and F4 broad peaks (’bumps’) in the spectrum, challenging. Manual inspection of these peaks is impractical due to the sheer quantity of segments requiring analysis.

**S2 Fig** shows the F1 and F2 range of rumbles within the six different events. Formants of different individuals within each event are marked by color. We notice that the formant ranges vary from event to event and we also see the difference in the range of formant frequencies among individuals. Here, again, all rumbles were sliced in segments of 0.25s before extracting the formant frequencies.

**S3 Fig** shows the narrowband spectrograms of the four rumlbles presented in Fig 4. The panels to the right of each spectrogram show the spectrum and LPC curve over, with F1 and F2 peaks, over the full duration of the rumble.

**S4 Fig** shows the formant trajectories in rumble exchange events 07a and 07b. These two events have a less prominent oscillatory behaviour, in particular event 07b which is rather flat, but is also the event with the least number of rumbles (eight in total).

**S5 Fig and S6 Fig** show the fit residuals for the two fits presented in Fig 9. The middle panel is the residuals plot and the bottom panel is the residuals vs fits plot.

**S7 Fig** shows an alternative fit for event 15, using a more complex sinusoidal function solution. The middle panel is the residuals plot and the bottom panel is the residuals vs fits plot.

An effort for a more detailed fit and residuals analysis is not of relevance at this stage, one reason being technical data considerations. More specifically, rumbles can sometimes overlap - with average overlap time of 1.5s [56]. Overlapping parts are removed (excised from the raw audio files) from our spectral analysis, as it is difficult to assign formant values in those sections. This means that in between some rumbles there are ‘missing’ segments. A more careful approach would require the separation of these overlapping signals using some type of source separation algorithm (something that we plan to do). An easier workaround would be to insert gaps equal to the duration of the removed overlapping sections and fit those data instead.

**S8 Fig - S12 Fig:** Five rumble exchange events are reordered in various ways, in order to examine how that would influence the presence of the observed oscillatory pattern presented in the section ‘Collectively oscillatory behaviour of rumbles’. In each figure the top panel shows the original event with the rumbles and segments of each rumble in their chronological order. In the second panel the order of the rumbles are randomized, but are grouped by the identity of the caller. In the third panel, the order of the rumbles is fully randomized. The fourth panel shows the rumbles in their original sequence, but the segments of each rumble are now scrambled. The last (bottom) panel shows a complete randomization of the sequence of all the segments (rumble slices) of the event. Two main interesting observations emerge from these figures: i) randomizing the rumble order can in some cases result in a new, different oscillatory pattern, ii) randomizing the order of segments within rumbles diffuses the oscillatory pattern observed in the original vocal exchange, indicative of the importance of the internal rumbles substructure in the construction of the oscillatory super-structure.

**S2 Fig.**
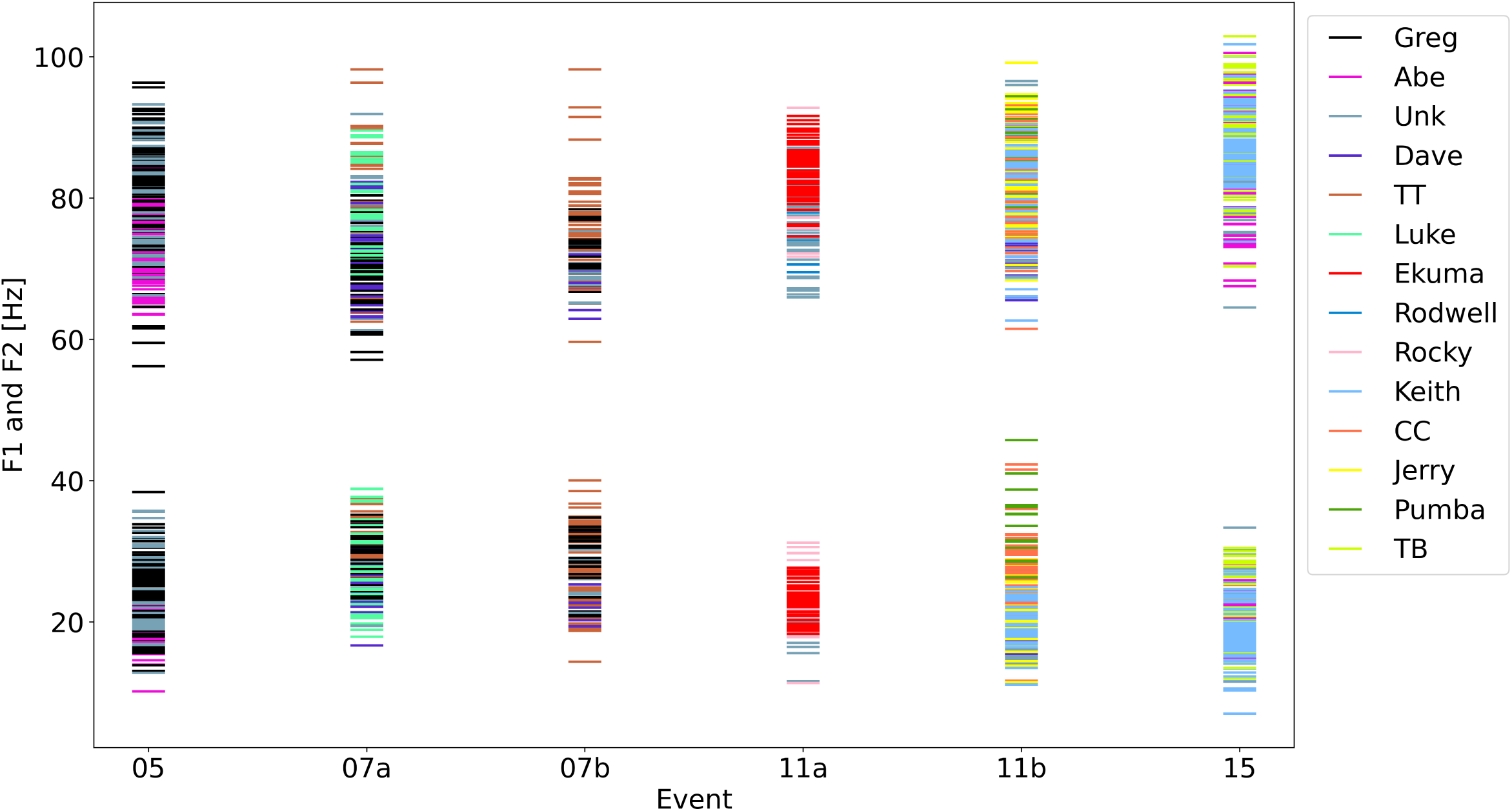
Formant frequencies F1 and F2 of rumbles within six different events. Formants of different individuals within each event are marked by color. We notice that the formant ranges vary from event to event. The difference in the range of formant frequencies among individuals is also visible. Each dash on the plot corresponds to a rumble segment of 0.25s.

**S3 Fig.**
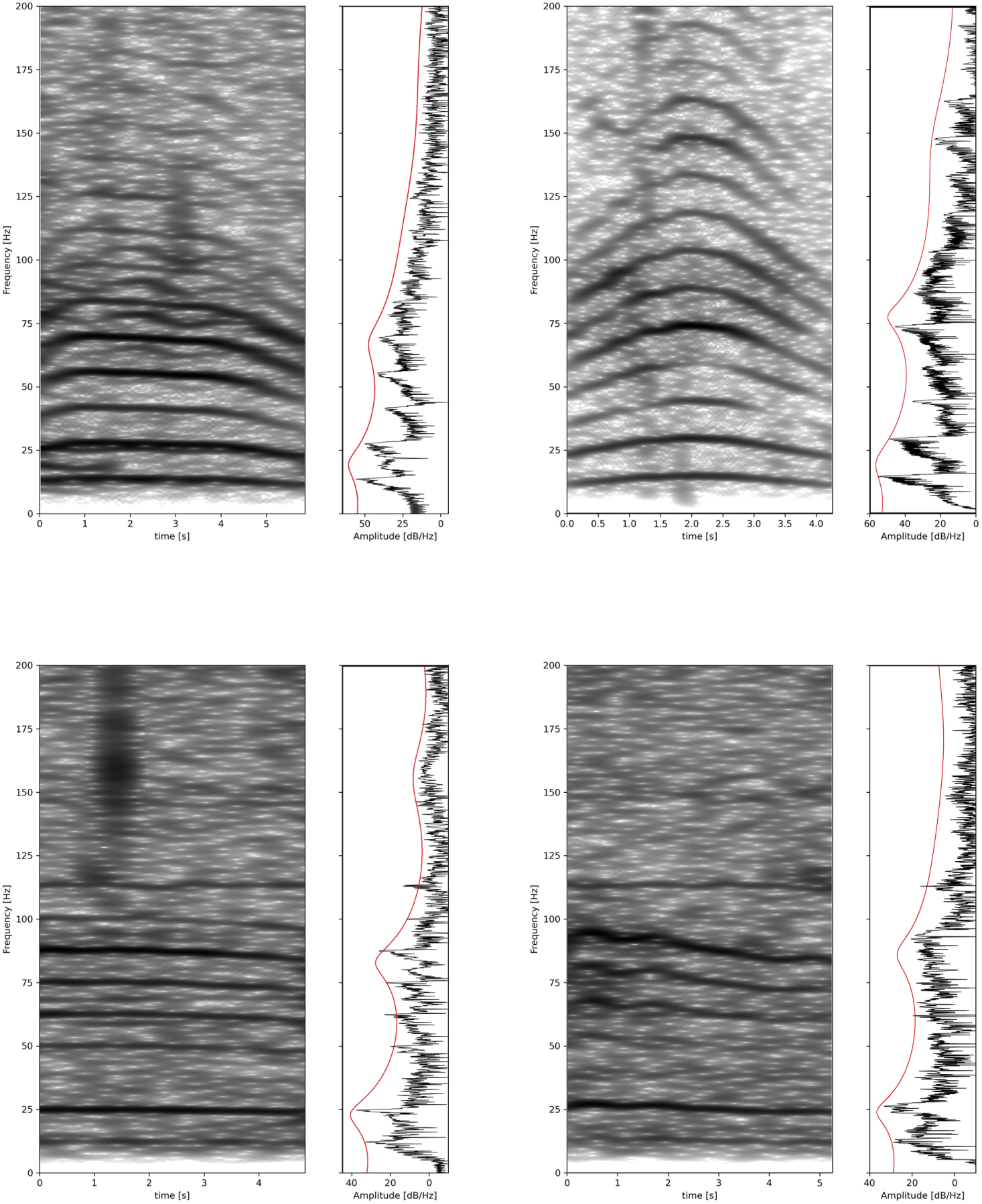
Narrowband spectrograms for four rumbles. The narrowband spectrograms for frequency range 0-200Hz of the rumbles presented in the broadband spectrograms of Fig 4 Each spectrogram is accompanied by its spectrum (black line in the window to the right) and the LPC curve in red showing the F1 and F2 ‘peaks’, this time calculated over the full rumble.

**S4 Fig.**
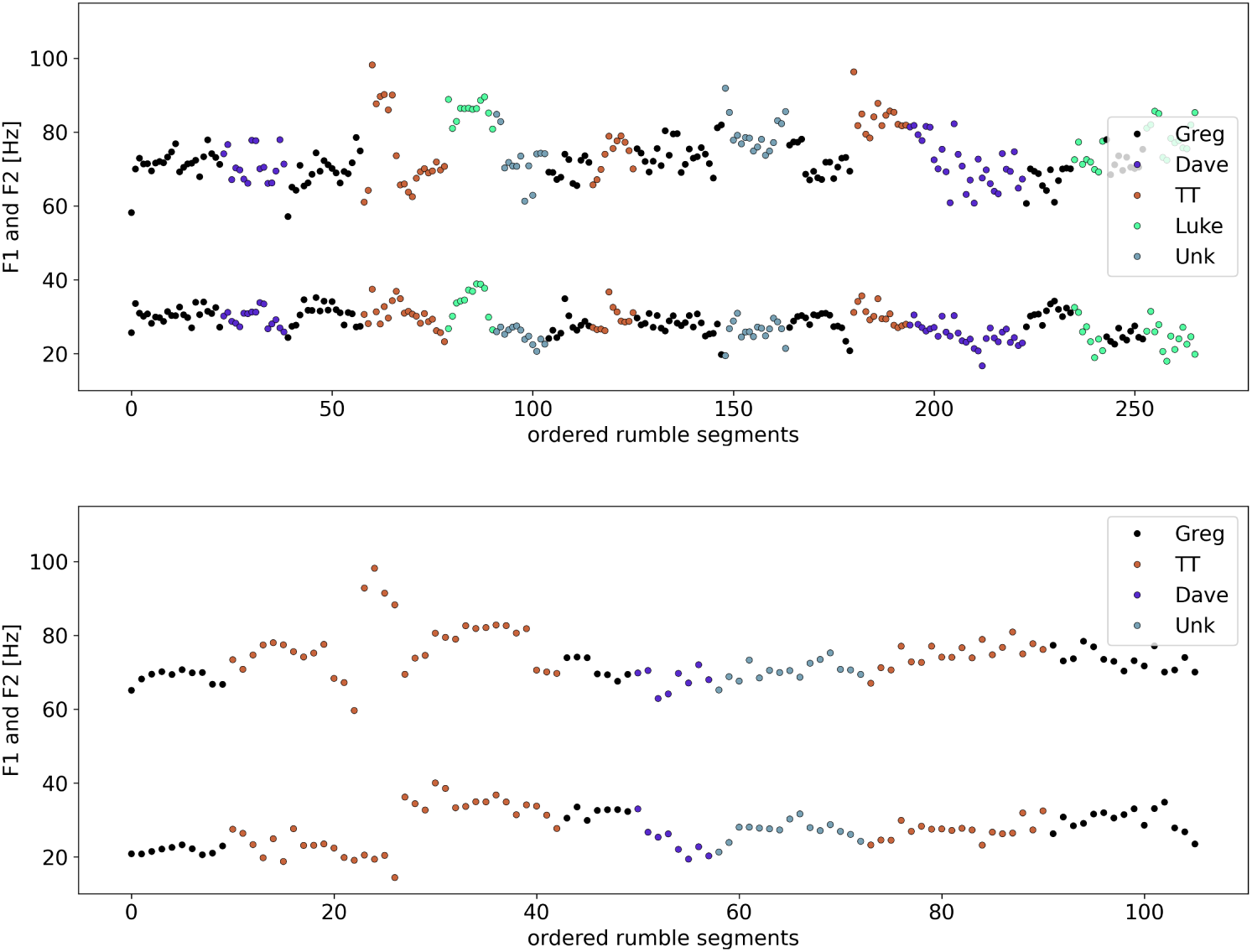
Formants trajectories in rumble exchange events 07a and 07b. The formants F1 and F2 of all rumbles in events 07a (top) and 07b (bottom) are plotted, retaining the chronological order of utterance of the rumbles. Each rumble was segmented in slices of 0.25s and each such slice is represented by a point. Color coding identifies the vocalizers. In these two events the oscillatory behaviour observed in the other events is not as prominent, although event 07a shows such behaviour in parts but not in its entirety. It’s notable that event 07b is the one with the fewest rumbles in our dataset.

**S5 Fig.**
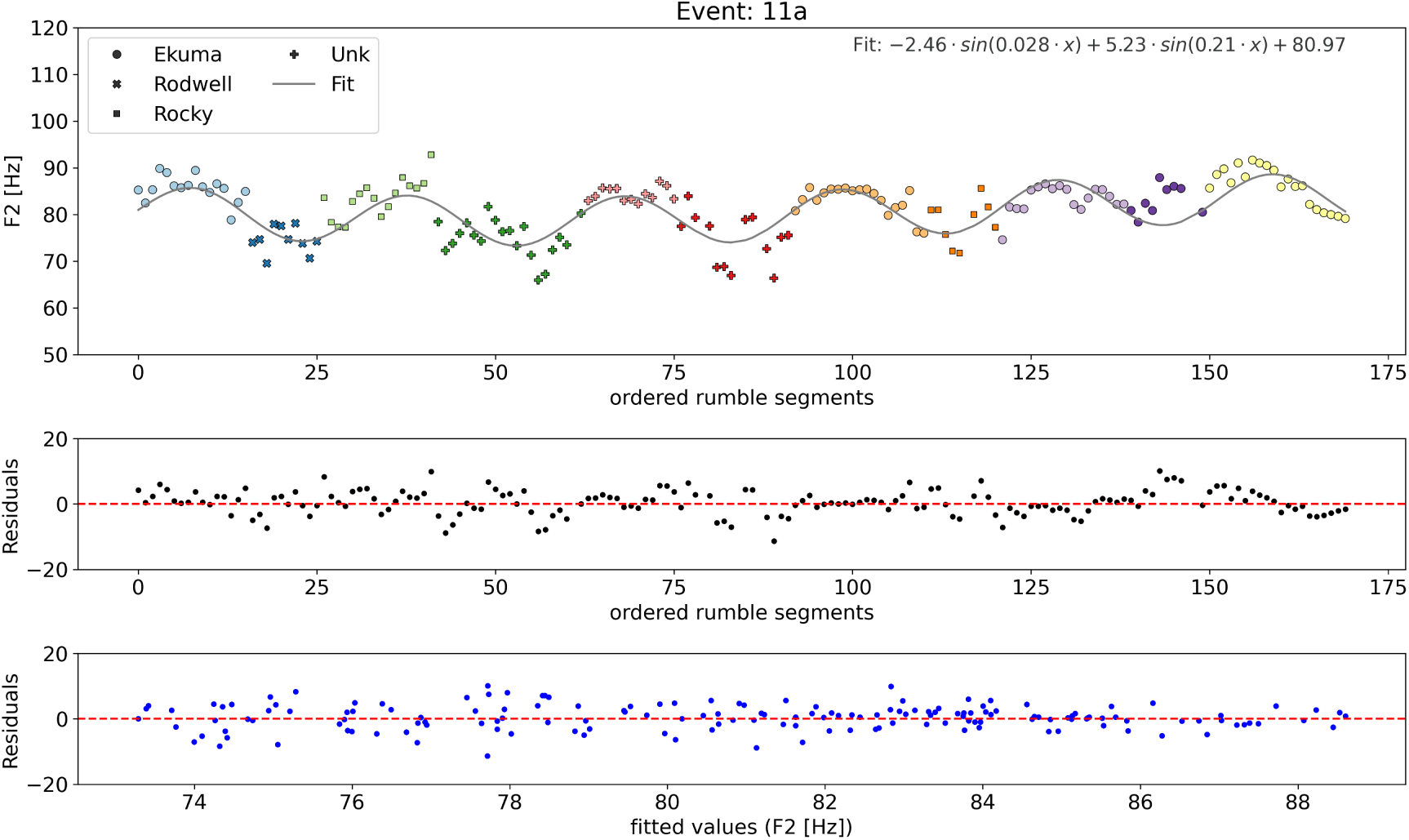
Fit and residuals for event 11a. The fit of event 11a as presented in Fig 9 with the residuals plot (middle) and the residuals vs fits plot (bottom).

**S6 Fig.**
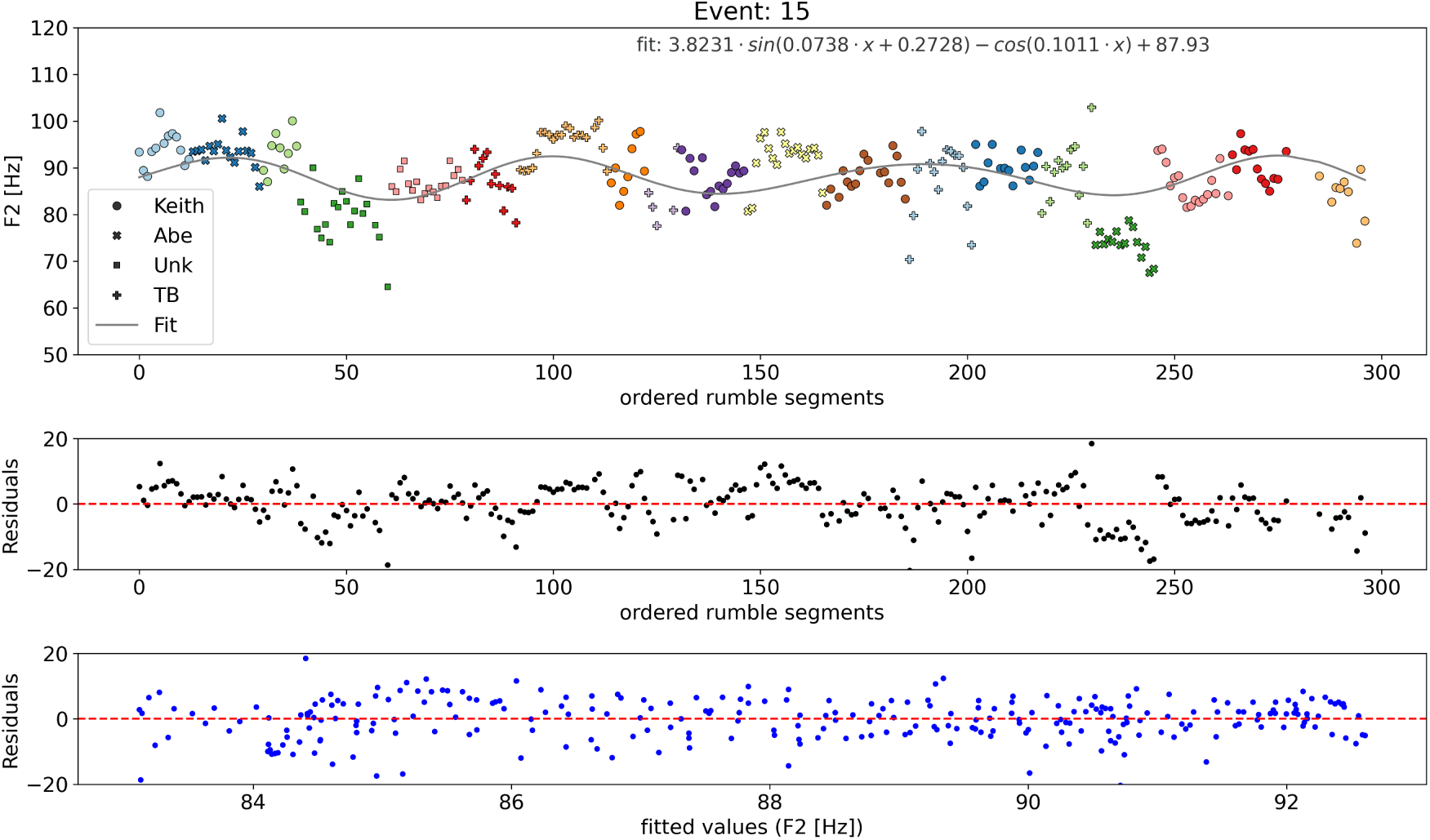
Fit and residuals for event 15. The fit of event 15 as presented in Fig 9 with the residuals plot (middle) and the residuals vs fits plot (bottom).

**S7 Fig.**
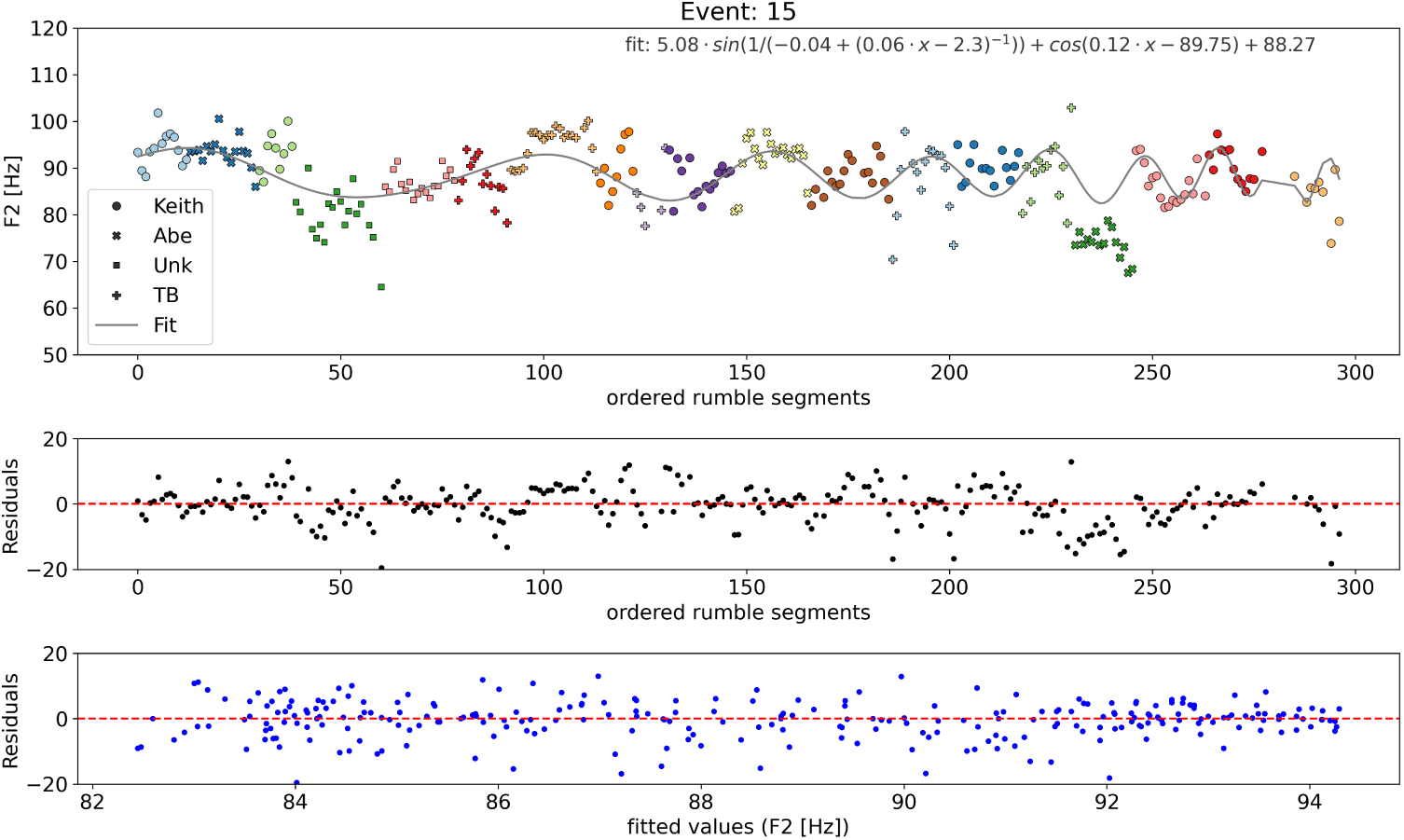
Alternative fit and residuals for event 15. A alternative, higher complexity fit of event 15, together with the residuals plot (middle) and the residuals vs fits plot (bottom).

**S8 Fig.**
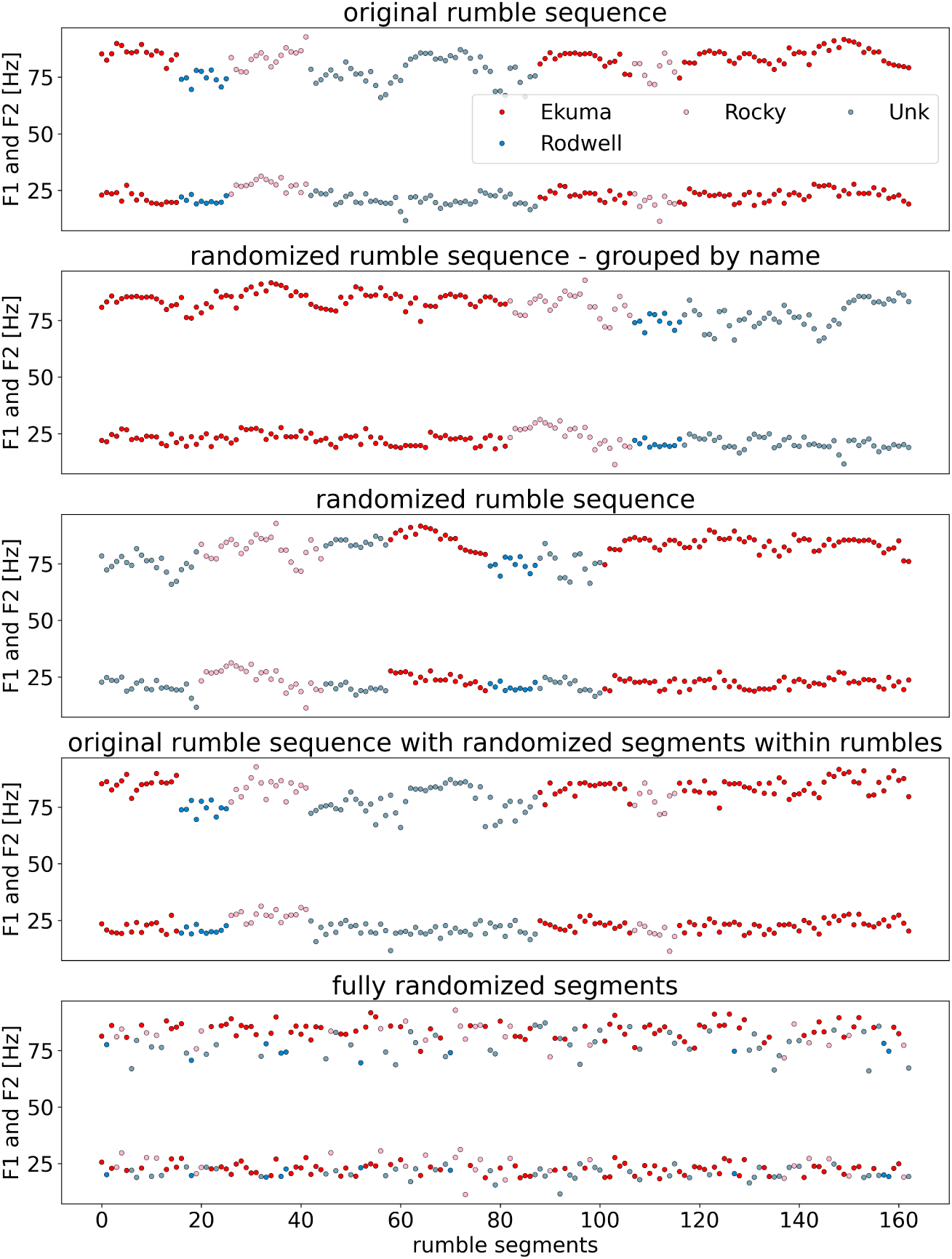
Scrambling event 11a. The order of rumbles within event 11a is randomly changed in various ways. From top to bottom: original order, randomly ordered rumbles but grouped by caller, randomly re-ordered rumbles within the event, original rumble order but randomly reordered segments within each rumble, complete randomization at the level of rumble segments.

**S9 Fig.**
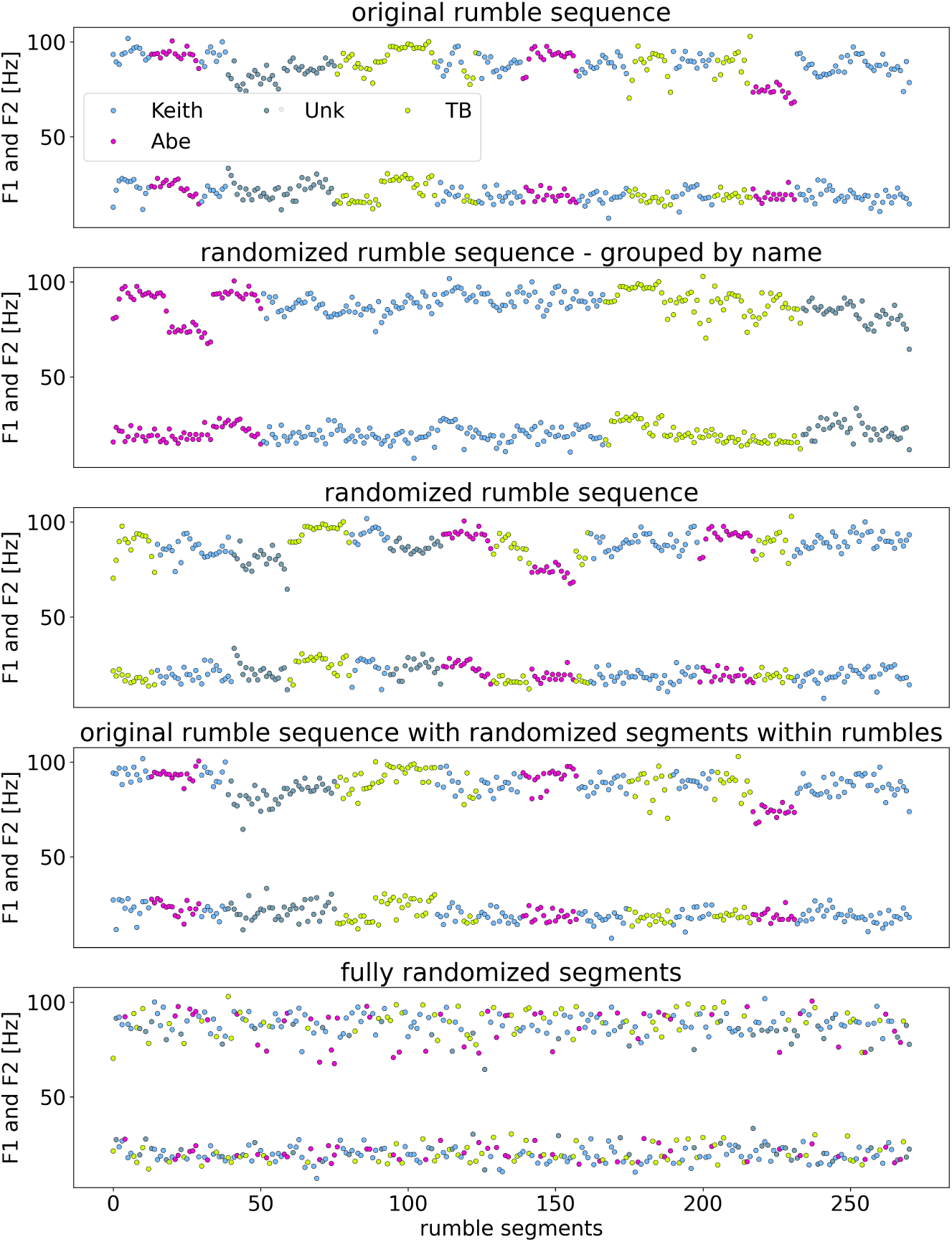
Scrambling event 15. The order of rumbles within event 15 is randomly changed in various ways. From top to bottom: original order, randomly ordered rumbles but grouped by caller, randomly re-ordered rumbles within the event, original rumble order but randomly reordered segments within each rumble, complete randomization at the level of rumble segments.

**S10 Fig.**
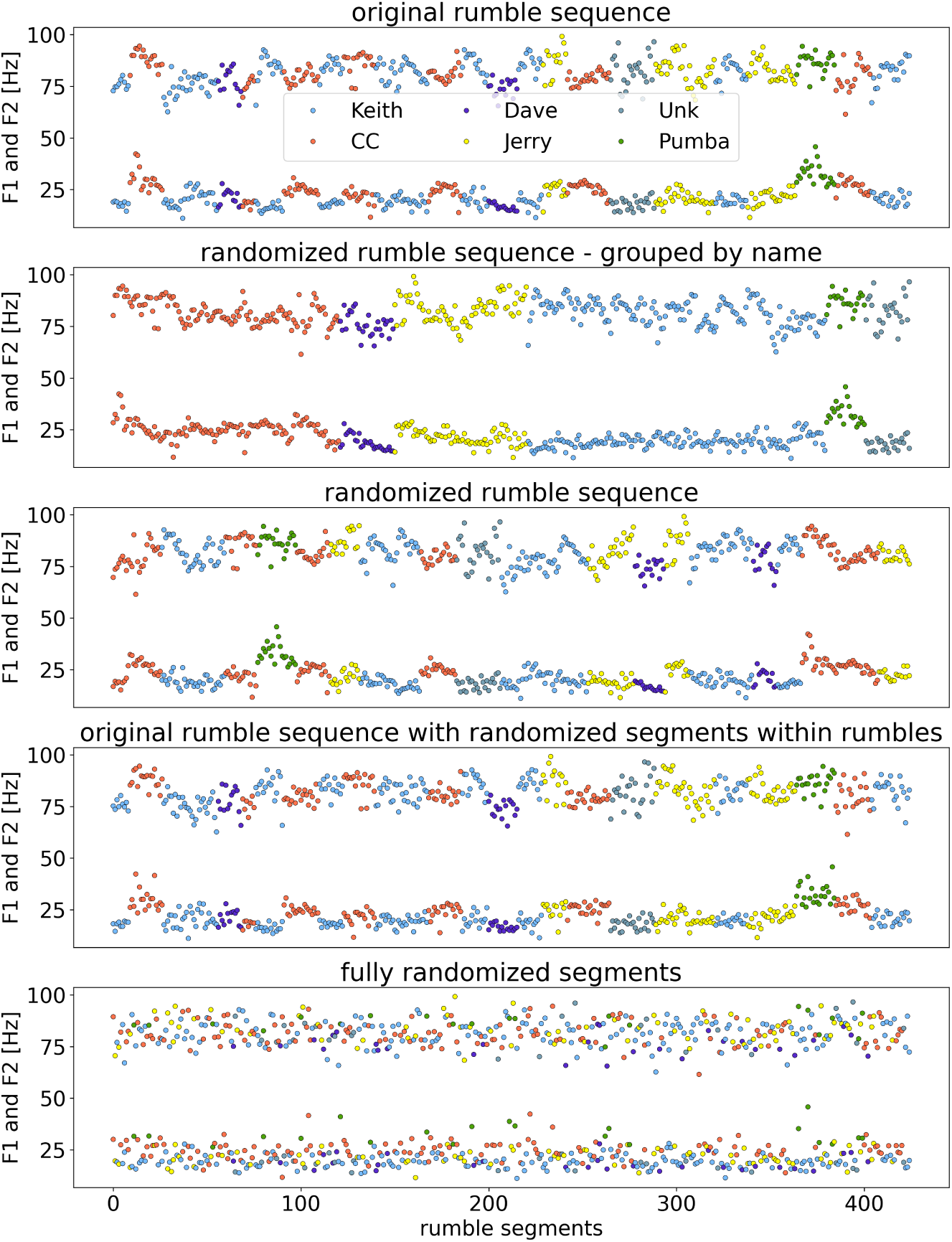
Scrambling event 11b. The order of rumbles within event 11b is randomly changed in various ways. From top to bottom: original order, randomly ordered rumbles but grouped by caller, randomly re-ordered rumbles within the event, original rumble order but randomly reordered segments within each rumble, complete randomization at the level of rumble segments.

**S11 Fig.**
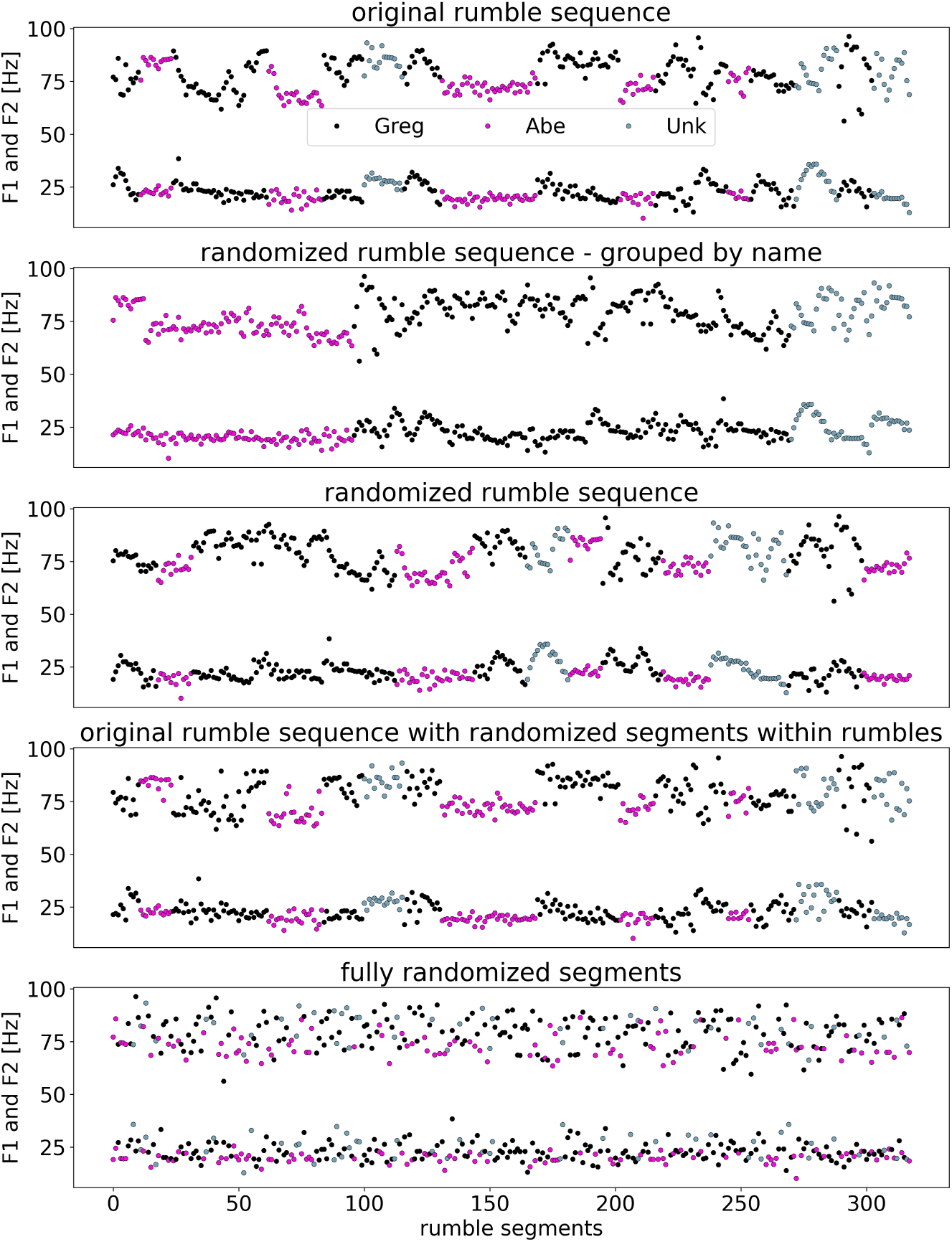
Scrambling event 05. The order of rumbles within event 05 is randomly changed in various ways. From top to bottom: original order, randomly ordered rumbles but grouped by caller, randomly re-ordered rumbles within the event, original rumble order but randomly reordered segments within each rumble, complete randomization at the level of rumble segments.

**S12 Fig.**
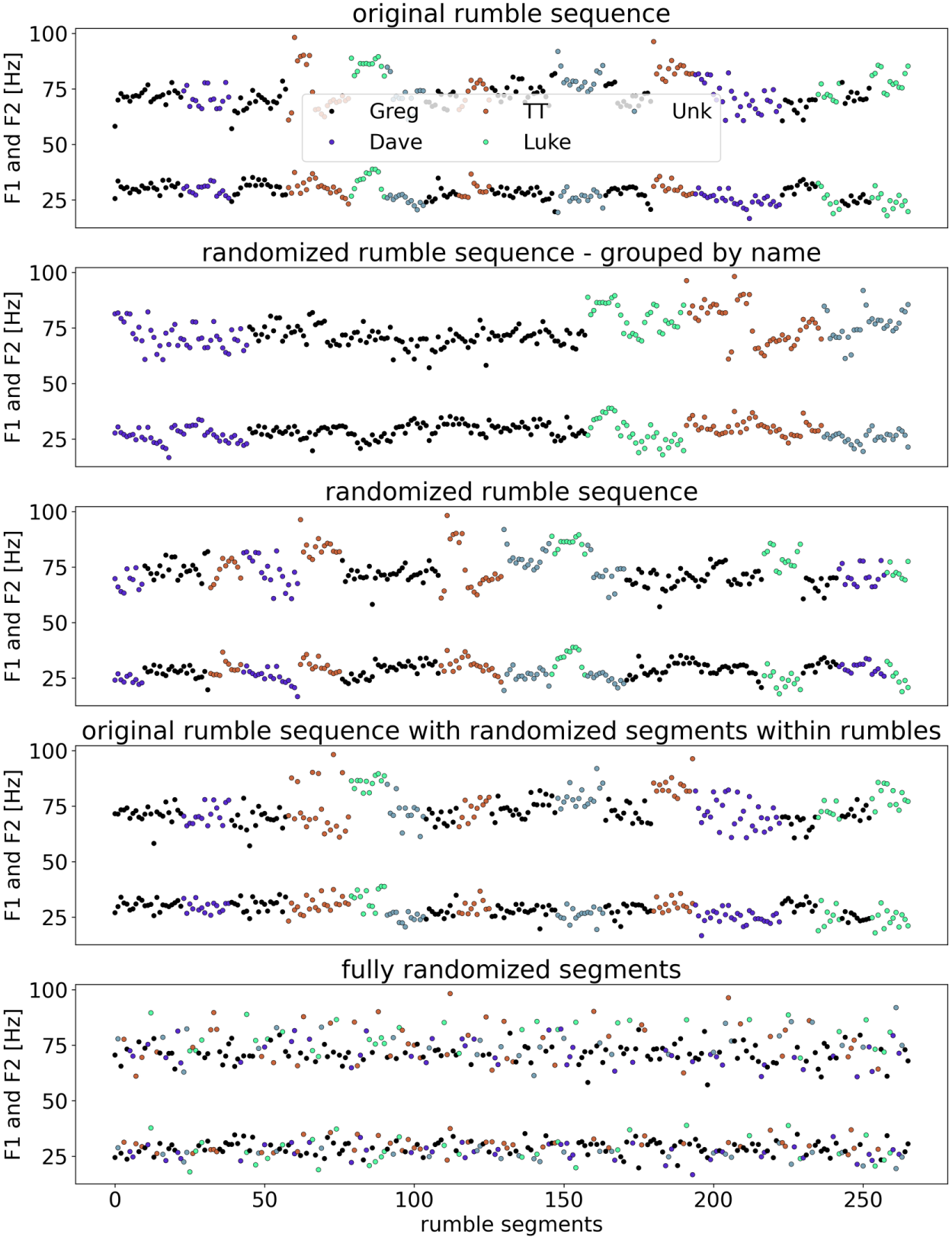
Scrambling event 07a. The order of rumbles within event 07a is randomly changed in various ways. From top to bottom: original order, randomly ordered rumbles but grouped by caller, randomly re-ordered rumbles within the event, original rumble order but randomly reordered segments within each rumble, complete randomization at the level of rumble segments.

