## Supplementary figures and images for "Elephant rumble vocalizations: spectral substructures and superstructures"

### Supp. Fig. 3

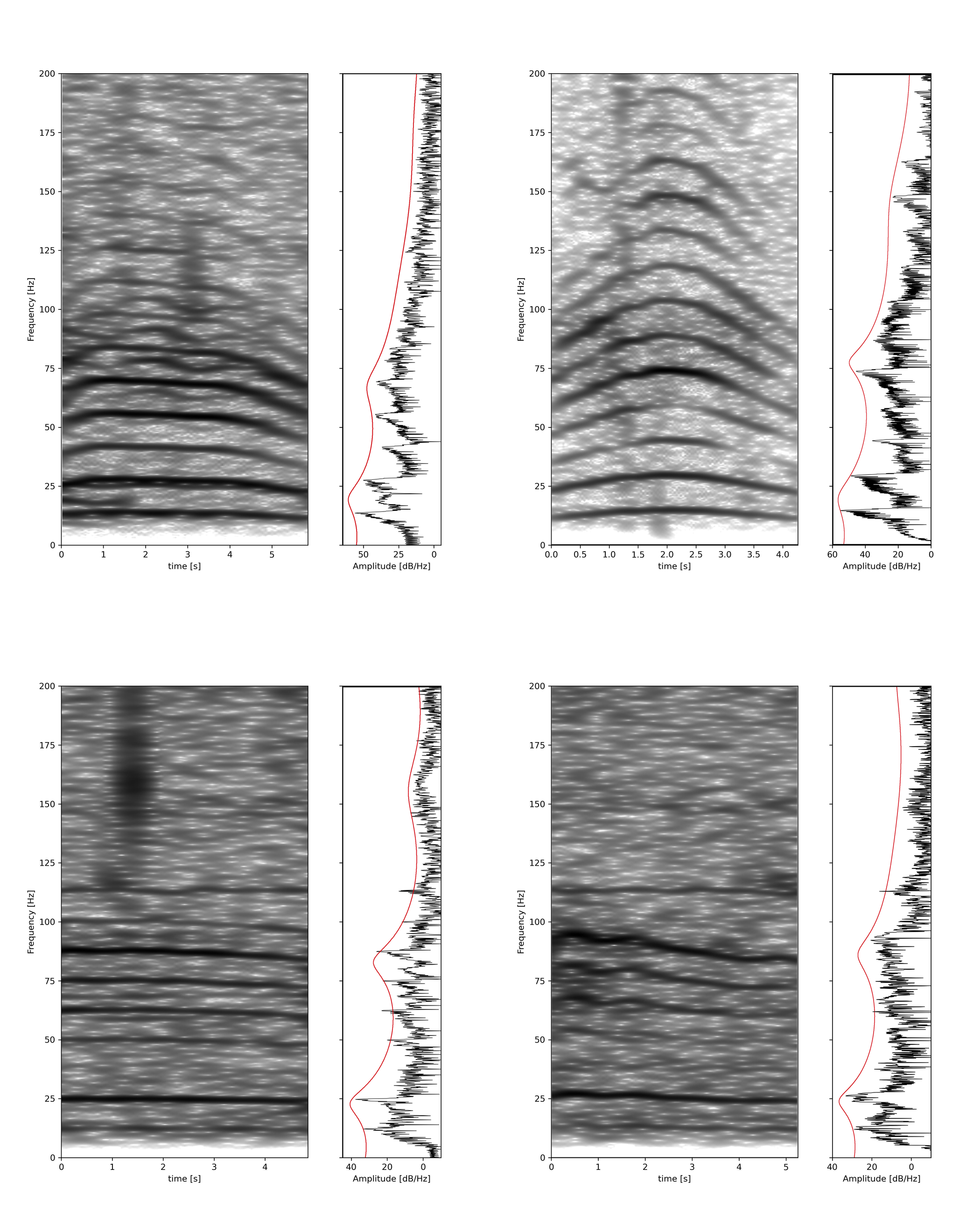
